# Lysosomal control of proteostasis and reproductive capacity by conserved LMD-3 protein in *C. elegans*

**DOI:** 10.1101/2025.05.16.654472

**Authors:** Yile Zhai, Tiantian Wang, Jiangxue Han, Mengxiao Wu, Meixi Gong, Wenfei Li, Zhe Zhang

**Author notes:** These authors contributed equally: Yile Zhai, Tiantian Wang. Correspondence (W. Li), (Z. Zhang).

## Abstract

The reproductive capacity declines markedly with female age in humans. Similarly, *C. elegans* exhibit a rapid age-dependent decline in fecundity shortly after reaching adulthood. Emerging evidence suggests a significant link between proteostasis disruption and reduced fertility in both worms and humans, but the regulatory mechanisms governing this connection are not fully understood. Here, we report that LMD-3, a LysM domain protein, regulates proteostasis and reproductive capacity in *C. elegans*. The deficiency of *lmd-3* leads to striking defects in oxidation resistance and constitutively high cellular stress responses, such as ER stress response and cytosolic stress response. We demonstrate that lysosome-localized LMD-3 protein interacts with vitellogenin and V-type ATPase, which drives proton-transporting and lysosomal lumen acidification. LMD-3 regulation of lysosomal function is essential for maintaining yolk protein homeostasis and reproductive health. We also identify that vitamin B12 (meCbl) supplementation revert the fecundity decline of *lmd-3* mutants by reducing oxidative stress and improving lysosomal function. Together, these findings emphasize the role of LMD-3 in sustaining protein homeostasis and oocyte quality control, and establish a model system to find potent therapeutic strategies to increase reproductive health.

## Introduction

Fecundity, the potential reproductive capacity, declines markedly with female age, which is one of the earliest aging phenotypes in many organisms, including yeast, worms and humans^1,2^. Societal impacts of declining fertility rates are profound, since the average maternal age at first childbirth has risen dramatically in modern society^3^. The risk of infertility, miscarriage and birth defects increases substantially during reproductive aging, with the decrease of oocyte quality being the major cause^4,5^. Although diverse factors, such as oxidative stress and mitochondrial dysfunction, have been suggested to contribute to reduced reproductive capacity^6^, the specific interplay between these factors and effective treatments remain elusive.

Because of the short life span, simple reproductive system and conserved genetic pathways, *Caenorhabditis elegans* is a prominent model for studying somatic aging and reproductive function^7,8^. Similar to human females, *C. elegan*s exhibits a long post-reproductive period and experiences a loss of reproductive capacity in mid-life, which is also caused by oocyte quality decline^9,10^. Recently, *C. elegans* research has substantially advanced our understanding of mechanisms that maintain reproductive capacity, and identified potential interventions to address age-related fertility decline^7,10,11^. In worms as in humans, maintaining proteostasis, such as protein folding, trafficking and degradation, is essential for oocyte quality and reproductive health^12,13^.

Oxidative stress can impair mitochondrial function and induce protein unfolding/misfolding, which further leads to proteostasis collapse^14^. Distinctively, oxidative stress in the ovaries deteriorates oocyte quality, accelerates oocyte and granulosa cell (GC) apoptosis, leading to a decline in the ovarian reserve^15^. Recently, Oxidation Resistance 1 (OXR1), a conserved but poorly characterized protein, has been shown to protect cells against oxidative damage in most eukaryotes, such as LMD-3 in *C. elegans* and Oxr1a in zebrafish^16–18^. OXR1 belongs to a group of protein that share a highly conserved TLDc domain, which interacts with V1 domain of V-ATPase and confers the neuroprotective function against oxidative stress^19^. Through its crucial function in antioxidant defense, whether OXR1 is involved in maintaining the proteostasis of oocyte and reproductive capacity remains unclear.

In this study, we characterize *lmd-3* that encodes a conserved LysM domain-containing protein and is essential for maintaining reproductive health in *C. elegans*. The deletion of *lmd-3* results in elevated oxidative stress and decreased brood size, the endpoint to evaluate reproductive capacity. Using CRISPR-mediated specific deletion of each *lmd-3* domains, we show that the carboxy termini of TLDc domain is essential for the physiological importance of LMD-3. From immunoprecipitation followed by mass spectrometry-based proteomic (IP-MS) analysis, we identified that LMD-3 interacts with vitellogenin and V-ATPase, which are primary egg yolk precursor and crucial component of the lysosomal proton pump, respectively. We find that LMD-3 deficiency leads to disrupted lysosomal function and the accumulation of vitellogenin, which consequently contributes to reproductive decline. Furthermore, methylcobalamin (meCbl), an active forms of vitamin B12, supplementation can ameliorate these defects, suggesting a potential therapeutic avenue for mitigating age-related fertility decline. These findings collectively elucidate the role of LMD-3 for lysosomal function and proteostasis in maintaining reproductive health, which may have implications for reproductive capacity decline in higher organisms.

## Results

### *lmd-3* loss leads to oxidative stress and mitochondrial dysfunction

The *C. elegans lmd-3* encodes a protein featuring LysM, GRAM, and TLDc domains, showing broadly evolutionarily conservation with the mouse and human OXR1 proteins (Supplementary Fig. 1a, b). To verify the oxidative stress phenotype, we used *C. elegans* mutants carrying the *tm2345* allele (Fig. 1a, Supplementary Fig. 2), a 619-bp deletion that disrupts the LMD-3 protein through a frameshift and severely reduces its expression (Fig. 1b, Supplementary Fig.3a, and Supplementary Table 1). These results indicate that *lmd-3(tm2345)* is a loss-of-function (LOF) strain.

**Fig 1.**
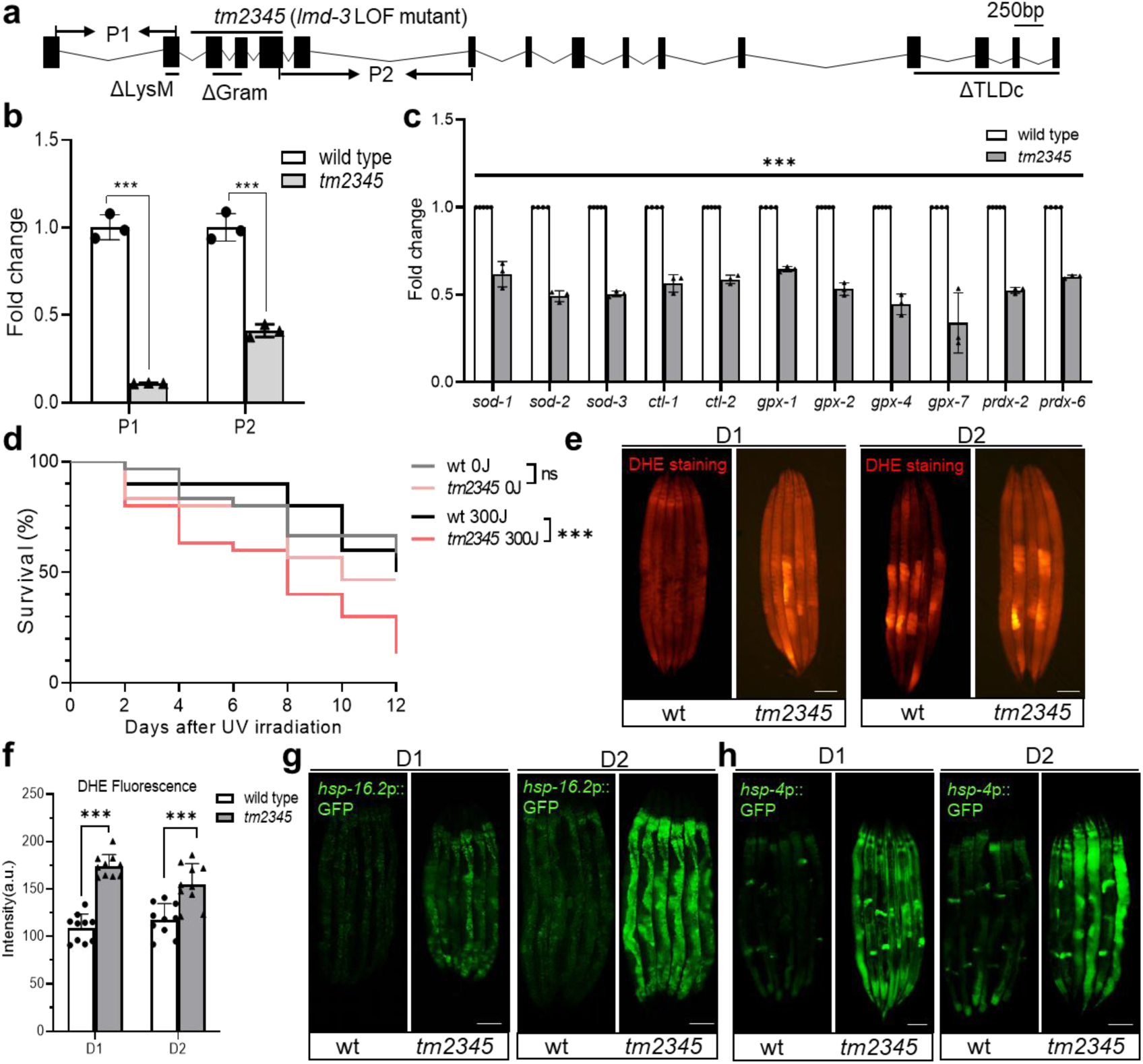
LMD-3 regulates ROS response in *C. elegans*. **a** Schematic of *lmd-3* LysM, GRAM, and TLDc domain deletion, and LOF mutant alleles with qRT-PCR P1 and P2 primers for *lmd-3* mRNA measurement. **b, c** qRT-PCR measurements of mRNA fold changes in wild-type and *tm2345* mutants (n = 3 for each group, unpaired t-tests: \*\*\**p* < 0.001) for endogenous *lmd-3* (**b**) and eleven selected antioxidant genes (**c**). **d** Survival curves of wild-type and *tm2345* mutant nematodes (n = 30 for each group, unpaired t-tests: \*\*\**p* < 0.001, ns, no significant differences) after UV irradiation (300 J/m^2^). **e, f** Superoxide anion (O₂⁻) level detection by DHE staining with exemplar images (**e**) and quantification (**f**) of DHE fluorescence in wild-type and *tm2345* (n = 14 for each group, unpaired t-tests: \*\*\**p* < 0.001). Scale bars, 100 μm. a.u., arbitrary units. **g, h** Exemplar fluorescence images for *hsp-16.2*p::GFP (**g**) and *hsp-4*p::GFP (**h**) with wild-type and *tm2345* mutants at D1 and D2 stage. Scale bars,100 μm. Source data are provided as a Source Data file.

We then assessed the expression of a panel of antioxidant genes in *tm2345* mutants, including superoxide dismutase (*sod-1, sod-2, sod-3*), catalase (*ctl-1, ctl-2*), glutathione peroxidase (*gpx-1, gpx-2, gpx-4, gpx-7*), peroxidase (*prdx-2*, *prdx-6*). Notably, the expression of all these genes was significantly reduced in *lmd-3(tm2345)* mutants (Fig. 1c and Supplementary Fig. 3b). In addition, when inducing reactive oxygen species (ROS) by exposing to 300 J/m² of UV irradiation^20^, *tm2345* mutants exhibited a significantly lower survival rate compared with wild type (Fig. 1d and Supplementary Fig. 3c), indicating increased susceptibility to oxidative stress. Consistently, using the fluorescent probe Dihydroethidium (DHE)^21^, we measured superoxide anion (O₂⁻) levels, precursors of ROS, in young adult animals, and found that *lmd-3(tm2345)* mutants displayed increased O₂⁻ accumulation (Fig. 1e, f).

As a consequence of elevated ROS, *tm2345* mutants exhibited a pronounced upregulation of *hsp-16.2*p::GFP (Fig. 1g and Supplementary Fig. 3d), an established cytoplasmic stress reporter in response to ROS-mediated damage^22,23^. Given that oxidative damage can disrupt ER structure and promote misfolded proteins aggregation, the loss of *lmd-3* also resulted a notable *hsp-4*p::GFP induction in *tm2345* mutants (Fig. 1h and Supplementary Fig. 3e), indicating ER stress and activation of the unfolded protein response (UPR)^24,25^. Furthermore, we tracked cytoplasmic stress levels throughout the *C. elegans* life cycle. Our results indicated that *tm2345* mutants experienced elevated cytoplasmic stress specifically during late developmental and early adult stages (Supplementary Fig. 3f, g).

To examine the potential role of *lmd-3* in mitochondrial protection, we analyzed mitochondrial morphology, number, and function in *tm2345* mutants. While *lmd-3* loss did not alter mitochondrial length and size (Supplementary Fig. 4a), but significantly reduced copy number and increased mtDNA damage in D1 adults (Supplementary Fig. 4b, c). Consistently, mitochondrial ATP levels and mitochondrial membrane potential were significantly reduced in *tm2345* mutants (Supplementary Fig. 4d-f). Taken together, these findings indicate essential roles of LMD-3 in antioxidant defense, maintaining cellular and mitochondrial homeostasis in *C. elegans*.

### Reduced fecundity and germ cell apoptosis induced by *lmd-3* deficiency in *C. elegans*

High levels of ROS and oxidative stress in the ovaries are known to cause premature ovarian failure (POF). This happens because they trigger apoptosis in oocytes and the surrounding granulosa cells (GCs), ultimately reducing the number of available eggs^26^. During our experiments, we consistently observed that *tm2345* mutants had fewer individuals than wild-type when grown under identical culturing conditions. This suggests that LMD-3 deficiency might lead to fewer offspring. Because reproductive processes are similar across many different organisms^8^, we’re now investigating exactly how LMD-3 deficiency affects reproduction and its control mechanisms in *C. elegans* (Supplementary Fig. 5).

Our analysis revealed a significant (20.4%) reduction in brood size of *lmd-3(tm2345)* mutants compared to wild-type animals across their reproductive lifespan, particularly evident during early reproductive stages (Fig. 2a, b and Supplementary Fig. 6a, b). Hypothesizing that this reduced fecundity might stem from germ cell deficiency^27^, we investigated the potential for *lmd-3* mutations to induce germ cell defects. The *C. elegans* germline, which producing sperm (stored in the spermatheca) and oocytes, possesses a distal stem cell region maintained by the distal tip cell (DTC), the primary component of the germline stem cell (GSC) niche. Germ cells proliferate distally before progressing proximally through meiotic prophase and later stages (Fig. 2c). This spatial arrangement facilitates the identification of meiotic stages by nuclear morphology and location^28^.

**Fig 2.**
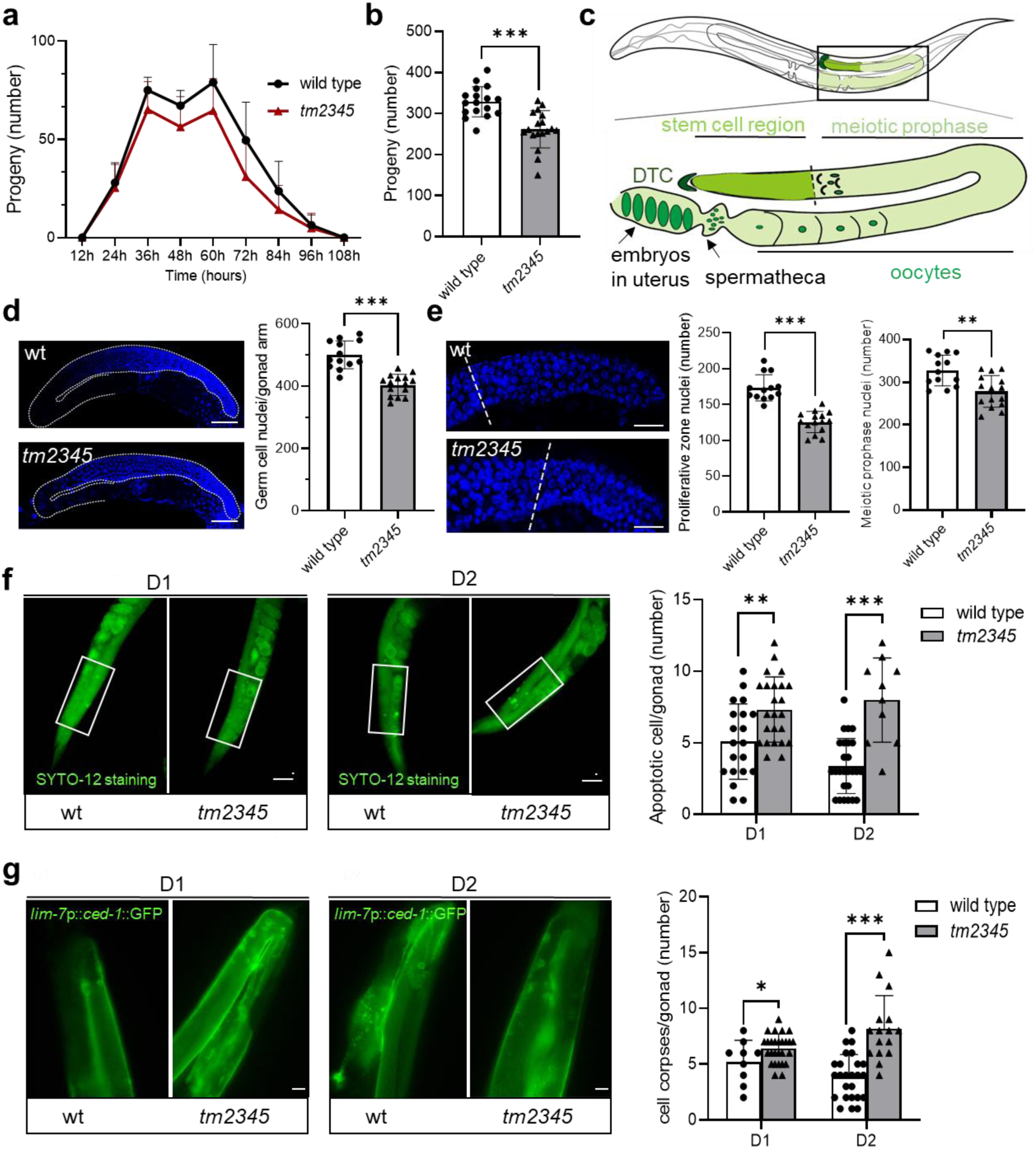
*lmd-3* deficiency induces germ cell apoptosis and reduces fecundity in *C. elegans.* **a** Progeny production curves of wild-type (black) versus *tm2345* (red) hermaphrodites that survive beyond their reproductive span. **b** Total brood sizes indicate the ability of *C*. *elegans* to produce offspring in wild-type and *tm2345* (n ≥ 13 for each group, unpaired t-tests: \*\*\**p* < 0.001). **c** *C. elegans* hermaphrodite germline consists of two distinct gonad arms, one illustrated in a schematic view: with a stem cell region and meiotic phase organized along the distal-proximal axis, which is ensheathed by the distal tip cell (DTC). **d** Exemplar fluorescence images and quantification of DAPI staining for total germline cells with wild-type and *tm2345* mutants (n ≥ 13 for each group, unpaired t-tests: \*\*\**p* < 0.001). Scale bars, 50 μm. **e** Exemplar fluorescence images and quantification of DAPI staining for germline stem cells and meiotic cells in wild-type and *tm2345* mutants (n ≥ 13 for each group, unpaired t-tests: \*\**p* < 0.01, \*\*\**p* < 0.001). Scale bars, 20 μm. **f, g** Exemplar fluorescence images and quantification for SYTO12 staining (**f**) and *lim-7*p::*ced-1*::GFP (**g**) with wild-type and *tm2345* mutants at indicated stage (n ≥ 9 for each group, unpaired t-tests: \**p* < 0.05, \*\**p* < 0.01, \*\*\**p* < 0.001). D1, Day 1 of adulthood. D2, Day 2 of adulthood. Scale bars, 50 μm. Source data are provided as a Source Data file.

To assess germ cell number, Day 1 adult gonads from wild-type and *tm2345* mutant animals were subjected to DAPI staining, a pan-nuclear DNA marker. Quantitative analysis of the resulting fluorescence signals demonstrated a pronounced decrease in the total number of germ cell nuclei in the mutant background (Fig. 2d, and Supplementary Fig. 6c). Furthermore, analysis of the germline stem cell region revealed a significant decline in the number of germline progenitor cells in the mutant strain at the same stage (Fig. 2e, and Supplementary Fig. 6d). These observations highlight the essential function of *lmd-3* in germ cell production and homeostasis, consistent with a fundamental germline developmental defect in its absence.

Considering LMD-3’s antioxidant defense and germ cell homeostasis roles in *C. elegans*, and the impact of oxidative stress on germline loss^29^, we investigated the role of apoptosis in the *tm2345* reduced germ cell phenotype. Two complementary methods were employed to evaluate germline apoptosis in *C. elegans*: SYTO12 staining, a nucleic acid dye exhibiting enhanced fluorescence in apoptotic cells^30^, and the *lim-7*p::*ced-1*::GFP reporter, allowing visualization of apoptotic bodies through green fluorescence^31^. Our assessment of apoptosis levels in *tm2345* mutants at both D1 and D2 adult stages clearly demonstrated a significant elevation in the number of apoptotic cells compared to wild-type animals (Fig. 2f-g, and Supplementary Fig. 6e-f). These findings establish that increased apoptotic activity is responsible for the reduced gonadal cell number in *lmd-3* mutants, consequently leading to a decreased number of germ cells and potentially contributing to a POF-like phenotype in *C. elegans*.

### LMD-3 mediates the reproductive capacity of *C. elegans* through its interaction with vitellogenin and V-ATPase proteins

To elucidate LMD-3’s role in the regulation of reproductive capacity, we generated transgenic expression of mCherry-tagged LMD-3 to identify its interacting proteins. Notably, only the N-terminal tagged LMD-3 was properly expressed, while the C-terminally tagged protein failed to accumulate to detectable levels (Fig. 3a). Likewise, abnormally elevated UPR monitored by *hsp-4*p::GFP was rescued by transgenic expression of N-terminally tagged *lmd-3* (Fig. 3b), but not the C-terminal form (Supplementary Fig. 7a). The C-terminal mCherry tag may impair LMD-3 expression and function by affecting protein folding or C-terminal TLDc domain integrity (Supplementary Fig. 1).

**Fig 3.**
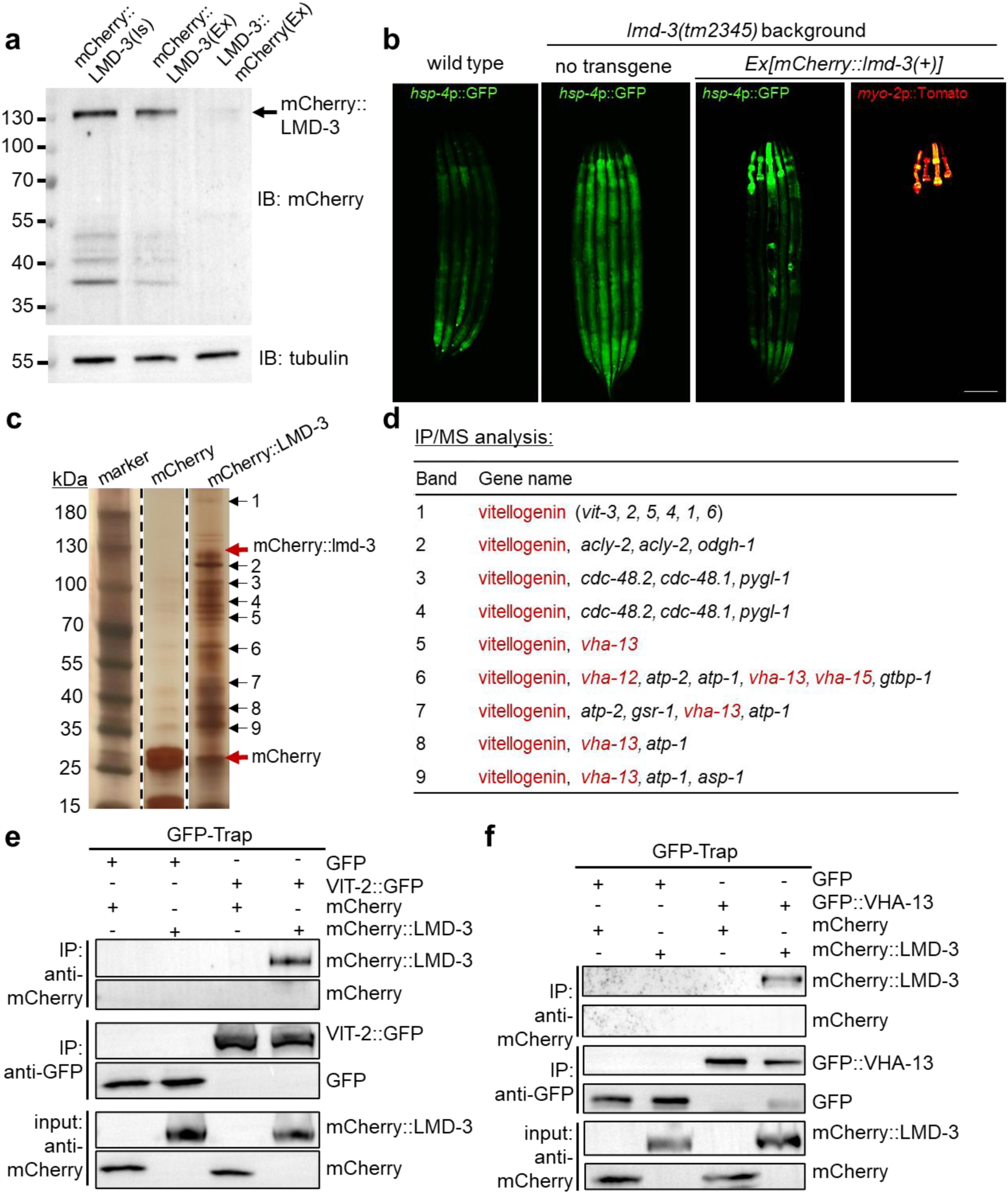
LMD-3 interacts with vitellogenins and V-ATPases to regulate cellular homeostasis. **a** Representative SDS-PAGE and Western blot analysis of transgenic LMD-3 expression from total animal lysates. LMD-3 was tagged with mCherry fluorescent protein at the N-terminus (N-term) and C-terminus (C-term). IB, immunoblotting; IS, integrated transgene; EX, extrachromosomal transgene. **B** Representative fluorescence images showing the rescue of *hsp-4*p::GFP induction in *lmd-3(tm2345)* mutants by ubiquitously expressed *rpl-28*p::mCherry::*lmd-3*. The trans gene is marked by pharyngeal *myo-2*p::mCherry. Scale bars, 50 μm. **c** Silver-stained SDS-PAGE showing affinity-purified proteins from whole worm lysate expressing mCherry::LMD-3 using RFP trap. Putative LMD-3-specific bands are indicated, and these bands were excised from the gel for mass spectrometry analysis. **d** Gene list of corresponding proteins identified in bands unique to LMD-3 (marked by numbers) is shown. Vitellogenin and V-ATPases coding genes are in red. **e, f** Co-Immunoprecipitation (co-IP) and western blot showing biochemical interaction of N- terminal mCherry-labeled LMD-3 with GFP-labeled VIT-2 (**e**) and GFP-labeled VHA-13 (**f**), respectively. Transgenic animals expressed with expression vectors, lysed for immunoprecipitation by GFP-TRAP, and blotted by antibodies against GFP and mCherry.

The utility of a functional mCherry::LMD-3 translational reporter, as evidenced by its rescue of the *tm2345* mutant phenotype (Fig. 3b), enabled the identification of LMD-3-interacting proteins. Affinity purification was performed from *C. elegans* lysates expressing this reporter using RFP-Trap magnetic beads, followed by silver-stained SDS-PAGE (Fig. 3c). Mass spectrometry (MS) analysis revealed vitellogenin proteins as the most abundant proteins identified (Fig. 3d). Vitellogenin, the principal precursor to yolk proteins essential for early embryonic development, is synthesized in somatic tissues and transported via the circulatory system to the developing oocytes across nearly all oviparous species^32^.

*C. elegans* possesses a family of six vitellogenin genes (*vit-1* to *vit-6*), whose expression is essential for both reproductive capacity and reproductive senescence^33,34^. The corresponding protein products of all six genes were identified as top-ranked in the analyzed gel slices by mass spectrometry (Supplementary Table 2). Of note, we also identified V-ATPase subunits (VHA-12, VHA-13, and VHA-15), established binding proteins of the TLDc domain of OXR1 in other species^35,36^, suggesting the validity of our mass spectrometry for profiling LMD-3 interactors (Fig. 3d, and Supplementary Table 2). V-ATPases acidify lysosomes for enzyme activity, essential for clearing germ cell corpse and oocyte protein aggregates, which maintains reproductive homeostasis^14^.

The interactions of LMD-3 with VIT-2 (Fig. 3e) and VHA-13 (Fig. 3f) were further substantiated through independent experimental validation (Supplementary Fig. 7b, c). Together, these results suggest that LMD-3 helps regulate cellular homeostasis and reproductive capacity by interacting with vitellogenin (for embryonic nutrition) and V-ATPases (for lysosomal function). Further analysis is warranted to fully elucidate the regulatory mechanisms of LMD-3, especially the TLDc domain.

### LMD-3 modulates lysosomal activity to ensure reproductive capacity in *C. elegans*

The interaction of LDM-3 TLDc domain with V-ATPases, integral lysosomal membrane proteins, prompted us to investigate LMD-3’s in vivo localization and the resulting lysosomal phenotypes in *lmd-3*-deficient mutants. Subcellular localization studies using a previously described transgenic reporter bearing a N-terminal fluorescent protein tag (Fig. 3a), revealed significant colocalization of mCherry::LMD-3 with the established lysosomal marker LMP-1::GFP (Fig. 4a, Supplementary Fig. 8a). In contrast, LMD-3 did not substantially colocalize with markers for other major organelles, including the endoplasmic reticulum, Golgi apparatus, or mitochondria (Supplementary Fig. 8b). Additionally, GFP::LMD-3 also exhibited high colocalization with additional lysosomal markers, NUC-1::mCherry and mCherry::Rab-7 (Supplementary Fig. 8c), collectively substantiating the primary lysosomal localization of LMD-3.

**Fig 4.**
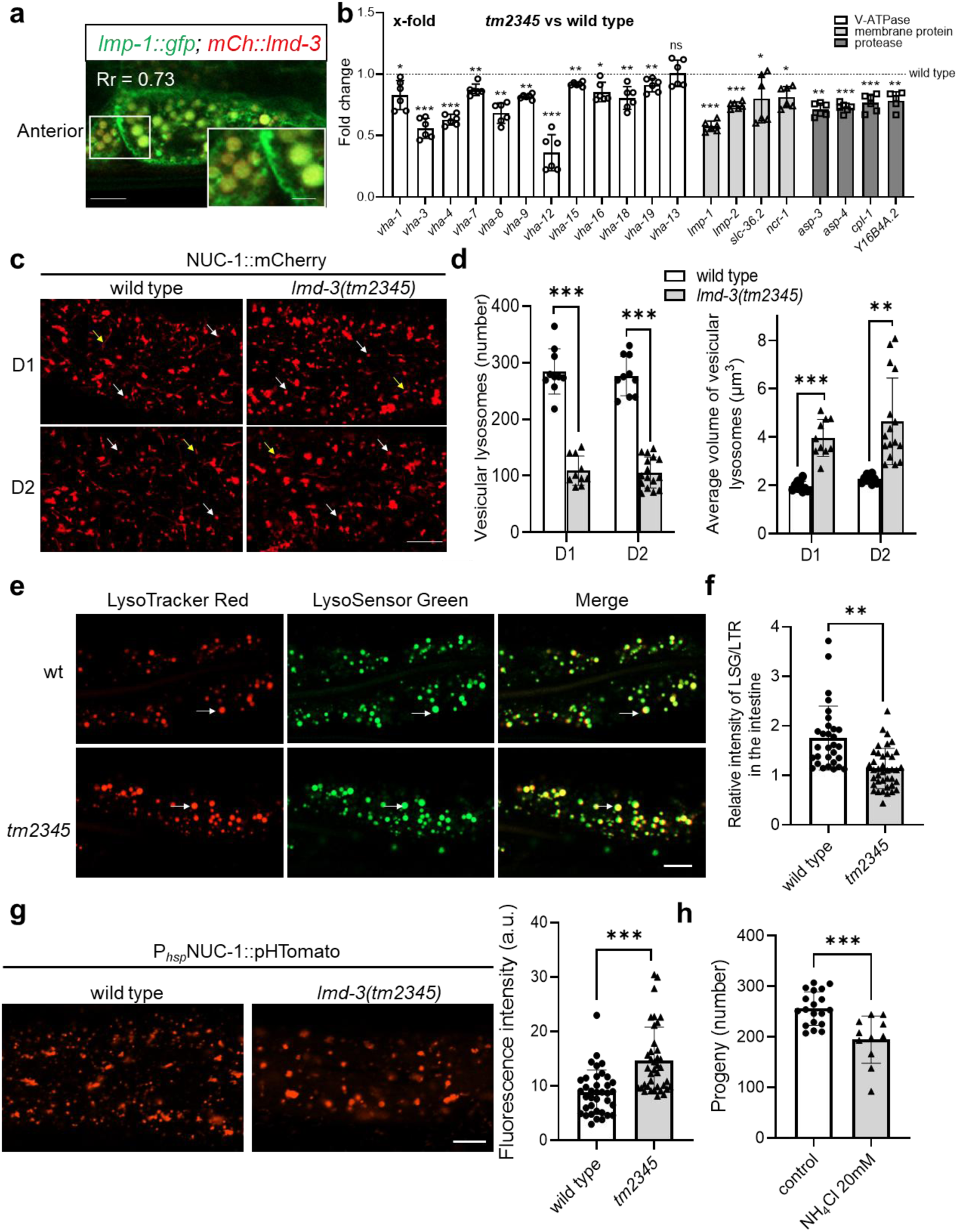
LMD-3 regulates lysosomal function and acidity in *C. elegans*. **a** Confocal image showing colocalization of mCherry::LMD-3 with the lysosomal marker LMP-1::GFP in the anterior region of the *C. elegans* hermaphrodite. Pearson’s correlation coefficient (Rr) indicates linear relationship of pixel intensities (Rr > 0.5 suggests good colocalization). Scale bar, 10 μm. **b** Transcriptional analyses in wild-type and *tm2345* animals at D1 stage (mean ± SEM, n=2 biological/3 technical replicates, unpaired t-tests: \**p* < 0.05, \*\**p* < 0.01, \*\*\**p* < 0.001, ns, no significant differences). **c, d** Confocal fluorescence images (**c**) and quantification (**d**) of the lysosomal reporter NUC-1::mCherry in wild-type and *lmd-3(tm2345)* animals at D1 and D2 adult stages. White and yellow arrows denote vesicular lysosomes and lysosomal tubules, respectively. Scale bar, 10 μm. Lysosome tubule number and volume were quantified (n ≥ 10 per group, unpaired t-tests: \*\**p* < 0.01, \*\*\**p* < 0.001). **e, f** Confocal fluorescence images (**e**) and quantification (**f**) of LSG DND-189/LTR DND-99 co-stained intestines of wild-type and *tm2345* animals at the D1 stage. White arrowheads indicate vesicular lysosomes stained by both LSG and LTR. Scale bars, 10 μm. Relative intensity of LSG/LTR were quantified (n ≥ 31 per group, unpaired t-tests: \*\**p* < 0.01). **g** Exemplar fluorescence images and quantification of the hypodermis at D1 adult, expressing NUC-1::pHTomato by the heat-shock (hs) promoter (n ≥ 37 per group, unpaired t-tests: \*\*\**p* < 0.001). a.u., arbitrary units. Scale bar, 10 μm. **h** Reproductive capacity of wild-type animals with 20 mM NH₄Cl treatment (n ≥ 11 per group, unpaired t-tests: \*\*\**p* < 0.001). Source data are provided as a Source Data file.

To evaluate the effect of *lmd-3* loss on the lysosomal machinery, we investigated transcriptional modulation of lysosome-related genes in *lmd-3(tm2345)* mutants. This analysis demonstrated a significant downregulation of the majority of transcripts encoding V-ATPase components (*vha* genes), lysosomal membrane proteins, and proteases (Fig. 4b, Source data 1). This finding indicates that LMD-3 is essential for maintaining the normal transcriptional output of a substantial portion of the lysosomal gene network.

We next examined lysosome morphology in wild-type and *lmd-3* mutants at D1 or D2 adult stage using NUC-1::mCherry reporter (Fig. 4c). Our quantitative analysis of demonstrated a reduced number of vesicular lysosomes and a significant increase in total lysosomal volume in *tm2345* mutants (Fig. 4d). These alterations suggest that *lmd-3* deficiency disrupts normal lysosomal dynamics and processing. Further investigation of lysosomal properties, including acidification and degradation activity, is needed to elucidate the underlying mechanisms.

Lysosome acidity was assessed by co-staining with LysoTracker Red DND-99 (LTR) and the acidotropic dye LysoSensor Green DND-189 (LSG, pKa 5.2), with LTR serving as a control for normalizing dye intake^37^. The quantified LSG/LTR fluorescence intensity ratio indicated lysosomal acidity, where increased LSG intensity reflects decreased pH (higher acidity). Our results revealed a decrease in lysosomal acidity in *lmd-3* mutants, suggesting decreased acidity (Fig. 4e, f). To further examine this, we employed a heat-shock inducible pH-sensitive reporter, NUC-1::pHTomato^38^. At 24 hours post-induction, this fluorescent protein localized to lysosomes, allowing intralysosomal pH assessment based on pHTomato’s increased fluorescence at higher pH (pKa ≈ 7.8)^38,39^. Corroborating the reduced LSG/LTR ratio, *tm2345* mutants exhibited significantly higher NUC-1::pHTomato fluorescence intensity compared to wild type (Fig. 4g). Collectively, these data indicate that LMD-3 modulates lysosomal acidity, likely through interaction with V-ATPases.

To elucidate the importance of lysosomal acidity for fertility in *C. elegans*, we aimed to neutralizing this acidity pharmacologically or genetically^40^. Treatment of wild-type animals with 20 mM NH_4_Cl, indeed significantly reduced their reproductive capacity (Fig. 4h). Similarly, genetic depletion of *vha-12*, a V-ATPase subunit interacting with LMD-3 (Fig. 3b, Supplementary Table 2), also decreased reproductive capacity compared to wild-type animals (Supplementary Fig. 9a). Furthermore, *lmd-3* mutants exhibited an additive reduction in reproductive capacity in the *vha-12(ok821)* background (Supplementary Fig. 9a), suggesting a synergistic requirement for LMD-3 and V-ATPases in lysosomal function essential for reproduction.

Given the interaction of LMD-3 with vitellogenin (Fig. 3c), and its role in lysosomal function and reproductive senescence^34^, we next assessed the potential link between LMD-3 and the regulation of lysosomal activity in vitellogenin processing. Using VIT-2::GFP, a reporter for this major vitellogenin, to study its expression and localization, we observed that *lmd-3* mutants exhibited an accumulation of VIT-2::GFP in the intestine, pseudocoelom, and germline (Supplementary Fig. 9b, c), suggesting impaired vitellogenin processing or uptake. While *vit-2* RNAi mildly increased lysosomal acidity in control animals, it failed to rescue the lysosomal acidity (Supplementary Fig. 9d) and reproductive defects (Supplementary Fig. 9e) of *tm2345* mutants. The potential functional redundancy among six vitellogenins in *C. elegans* might explain this lack of rescue.

To further assess LMD-3’s role in lysosomal degradation, we examined the clearance of secreted ssGFP, driven by the muscle-specific *myo-3* promoter and endocytosed by coelomocytes from the body cavity^41^. In line with the observed impairment in lysosomal acidity, *tm2345* mutants showed increased ssGFP fluorescence in coelomocytes (Supplementary Fig. 10a, b), indicating reduced lysosomal degradation. Further supporting this, *tm2345* mutants also exhibited enhanced aggregation of the polyglutamin proteostasis reporter *unc-54*p::Q40::YFP (Supplementary Fig. 10c, d)^42^, suggesting disrupted protein homeostasis in *lmd-3* mutants. These results indicate that LMD-3 deficiency causes lysosomal dysfunction, impacting reproduction and proteostasis, potentially linked to its interaction with V-ATPases and vitellogenin processing.

### TLDc domain of LMD-3 is essential for lysosomal function and reproductive capacity in *C. elegans*

The OXR1 protein family features conserved LysM (peptidoglycan binding)^43^, GRAM (lipid binding)^44^, and TLDc (V-ATPase interaction)^19^ domains (Supplementary Fig. 1). To elucidate the specific roles of LMD-3’s domains in regulating cellular homeostasis and reproductive capacity in *C. elegans*, we generated precise deletion mutants using CRISPR-Cas9 for the LysM (Δ22-67), GRAM (Δ135-203), and TLDc (Δ627-835) domains, respectively (Fig. 1a).

Notably, the lysosomal degradation defect observed in *tm2345* animals, characterized by strong *myo-3*p::ssGFP aggregation in coelomocytes, was specifically phenocopied by the TLDc deletion mutant (ΔTLDc) (Fig. 5a, b). Transcriptional analysis, however, revealed that unlike the *tm2345* frameshift allele, the TLDc deletion did not alter the overall *lmd-3* mRNA level (Supplementary Fig. 11a, b). Furthermore, cycloheximide (CHX, a ribosome inhibitor) chase assays demonstrated accelerated degradation of only ΔTLDc and *lmd-3(tm2345)* proteins (Supplementary Fig. 11c, d), indicating the TLDc domain’s essential role in LMD-3 protein stability.

**Fig 5.**
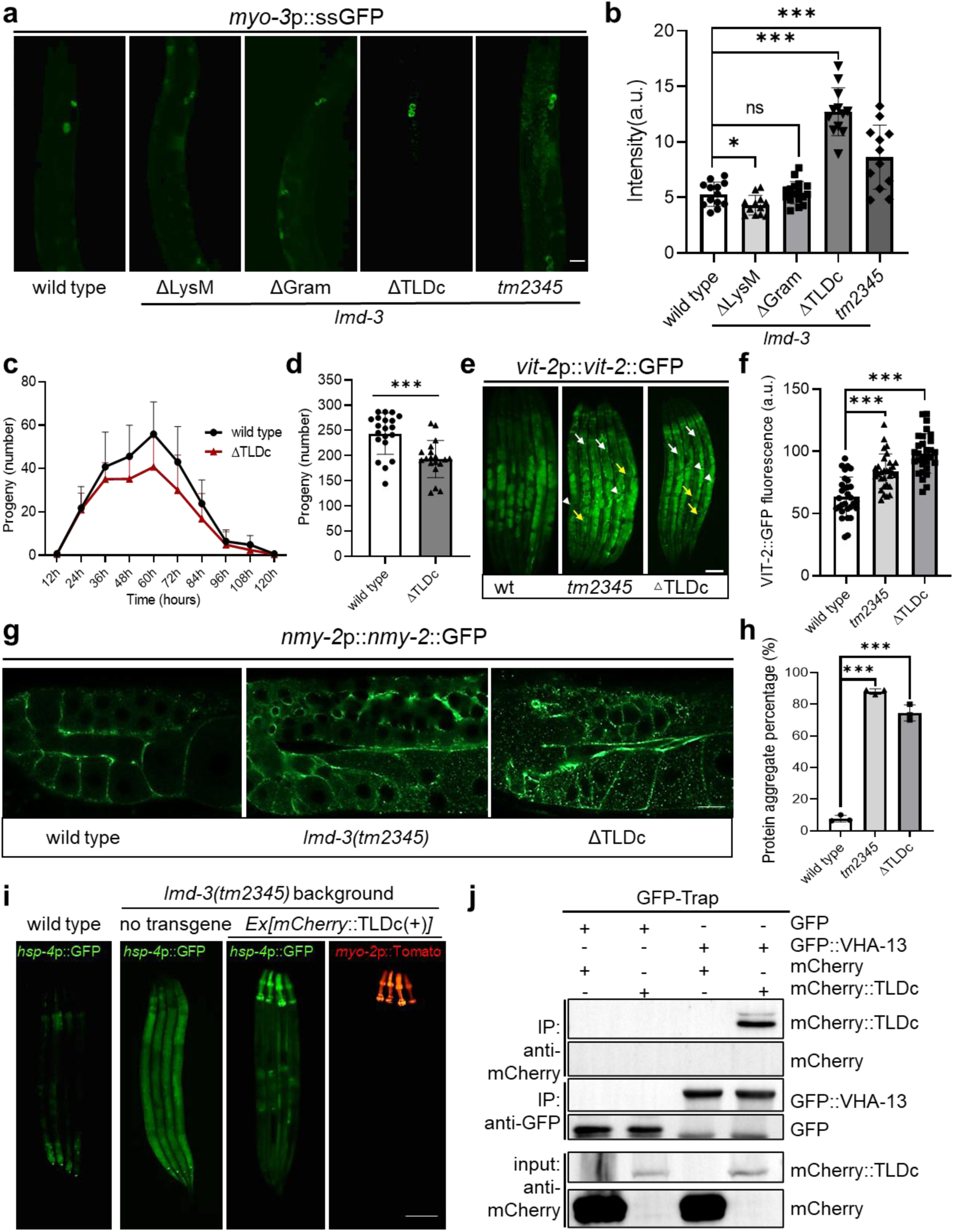
The role of the TLDc domain in LMD-3’s function. **a,b** Exemplar fluorescence images (**a**) and quantification (**b**) *myo-3*p::ssGFP accumulation in coelomocytes of wild-type and *lmd-3* mutant animals at D1 stage (n ≥ 12 for each group, unpaired t-tests: \**p* < 0.05, \*\*\**p* < 0.001, ns, no significant differences). Scale bars, 50 μm. **C** Progeny production curves of wild-type (black) versus TLDc deletion (red) hermaphrodites that survive beyond their reproductive span. **d** Total brood sizes indicate the ability of *C*. *elegans* to produce offspring in wild-type and ΔTLDc mutants (n ≥ 19 for each group, unpaired t-tests: \*\*\**p* < 0.001). **e, f** Exemplar fluorescence images (**e**) and quantification (**f**) *vit-2*p::*vit-2*::GFP accumulation in wild-type, *tm2345* and ΔTLDc mutants at D2 stage (n ≥ 30 for each group, unpaired t-tests: \*\*\**p* < 0.001). Aggregates in different tissues: white arrowheads denote germline, white arrows denote pseudocoelom, yellow arrows denote intestines. a.u., arbitrary units. Scale bars, 50 μm. **g, h** Confocal fluorescence images (**g**) and quantification (**h**) *nmy-2*p::*nmy-2*::GFP accumulation in wild-type, *tm2345* and ΔTLDc mutants at D1 stage(mean ± SEM, n=3 biological replicates, animals per condition per replicate = 30, unpaired t-tests: \*\*\**p* < 0.001). **i** Exemplar fluorescence images showing the rescue of *hsp-4*p::GFP induction in *lmd-3(tm2345)* mutants with transgenic expression of *lmd-3*p::TLDc::mCherry marked by pharyngeal *myo-2*p::mCherry. Scale bars, 50 μm. **j** Biochemical interaction of GFP-VHA-13 and mCherry-TLDc fragment demonstrated by co-IP and western blot of transgenic animal cell lysates. Co-transfected lysates were immunoprecipitated with GFP-Trap and immunoblotted for GFP and mCherry. Source data are provided as a Source Data file.

Consistent with the reproductive defect observed in *tm2345* animals, ΔTLDc mutants also exhibited reduced reproductive ability (Fig. 5c, d). Mechanistically, similar to the *lmd-3* null allele, ΔTLDc mutants displayed significantly increased ROS levels (Supplementary Fig. 11e-g), abnormally accumulated yolk proteins in the intestine, pseudocoelom, and germline (Fig. 5e, f), potentially contributing to lysosomal dysfunction. To monitor this, we utilized the heat-shock inducible lysosomal reporter P*hsp*NUC-1::sfGFP::mCherry^45^. The increased yellow puncta observed in *tm2345* and ΔTLDc animals (Supplementary Fig. 11h, i) indicate impaired lysosomal acidification and maturation, where GFP is quenched in acidic lysosomes, leaving only mCherry in mature, acidic structures.

Additionally, both ΔTLDc and *lmd-3(tm2345)* mutants showed increased aggregation of the age-related protein NMY-2::GFP in oocytes (Fig. 5g, h), further compromising oocyte protein homeostasis and likely contributing to reduced fertility. Significantly, overexpression of the LMD-3 TLDc domain alone rescued the *hsp-4p::GFP* upregulation in *lmd-3* mutants (Fig. 5i), and co-immunoprecipitation confirmed a conserved interaction between the TLDc domain and V-ATPase subunits (Fig. 5j, Supplementary Fig. 11j). The conserved TLDc domain of LMD-3 is crucial for cellular homeostasis, lysosomal function, protein stability, and reproduction in *C. elegans*, potentially via V-ATPase interaction and preventing yolk protein/oxidative stress accumulation.

### Vitamin B12 supplementation rescues reproductive defects in *lmd-3* mutants by alleviating cellular stress and improving organelle function

As *lmd-3* dysfunction impairs lysosomal function and reproductive health, accompanied by increased cellular stress, we explored whether enhancing stress resistance could restore fertility in *lmd-3* mutants. The *E. coli* K-12 strain HT115 has recently been shown to enhance *C. elegans* resistance to various cellular stresses and mitigate health defects associated with the standard OP50 (*E. coli* B strain) diet^46,47^. Based on this, we observed a complete rescue of the elevated cytoplasmic stress reporter *hsp-16.2*p::GFP in *tm2345* larvae transferred to an HT115 diet, compared to those fed OP50 (Fig. 6a). Importantly, culturing *lmd-3* mutants on the HT115 diet fully restored their reproductive ability (Fig. 6b) and germline morphology (Supplementary Fig. 12a, b). Together, these results indicate that reducing cellular stress via the HT115 diet can overcome the reproductive and developmental defects caused by *lmd-3* dysfunction.

**Fig 6.**
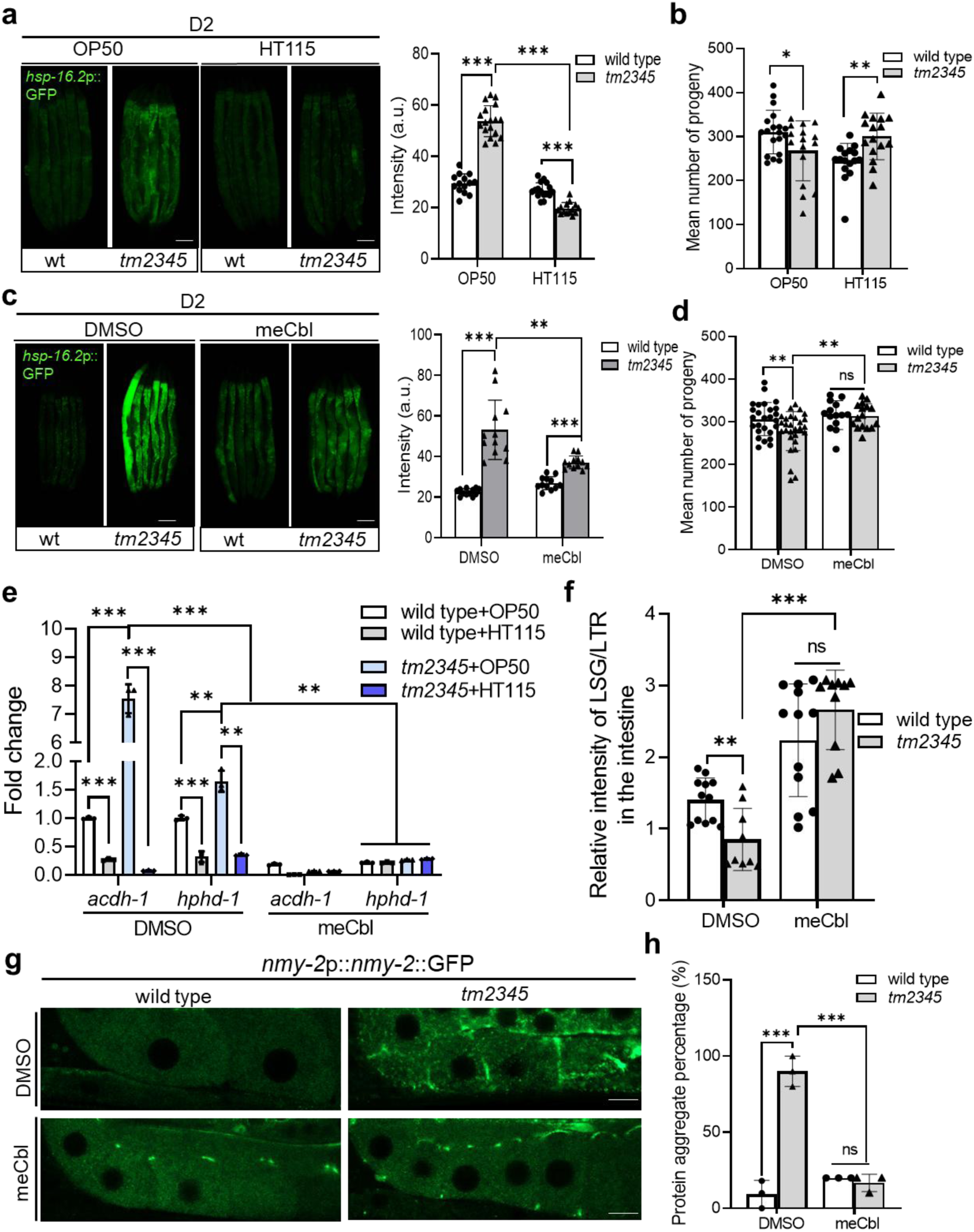
Vitamin B12 supplementation alleviates cellular stress and restores reproductive function in *lmd-3* mutants. **a** Exemplar fluorescence images and quantification *hsp-16.2*p::GFP of wild-type and *lmd-3* mutant animals feeding with OP50 or HT115 at D2 stage (n ≥ 14 for each group, unpaired t-tests: \*\*\**p* < 0.001). Scale bars, 100 μm. **b** Total brood sizes indicate the effect of H115 feeding on reproductive capacity in wild-type and *tm2345* mutants (n ≥ 16 for each group, unpaired t-tests: \**p* < 0.05, \*\**p* < 0.01). **c** Exemplar fluorescence images and quantification *hsp-16.2*p::GFP of wild-type and *lmd-3* mutant animals feeding with 0.15 μM meCbl at D2 stage (n ≥ 12 for each group, unpaired t-tests: \*\**p* < 0.01, \*\*\**p* < 0.001). Scale bars, 100 μm. **d** Total brood sizes indicate the effect of 0.15 μM meCbl supplementation on reproductive capacity in wild-type and *tm2345* mutants (n ≥ 14 for each group, unpaired t-tests: **P < 0.01, ns, no significant differences). **e** qRT-PCR measurements of mRNA fold changes in wild-type and *tm2345* mutants for *acdh-1* and *hphd-1* genes in different treatment conditions (n = 3 for each group, unpaired t-tests: \*\**p* < 0.01, \*\*\**p* < 0.001). **f** Quantification lysosomal acidity with LSG/LTR co-stained intestines of meCbl-treated wild type and *lmd-3(tm2345)* animals at the D1 stage (n ≥ 36 for each group, unpaired t-tests: \*\**p* < 0.01, \*\*\**p* < 0.001, ns, no significant differences). Scale bars. 10 μm. **g, h** Confocal fluorescence images (**g**) and quantification (**h**) *nmy-2*p::*nmy-2*::GFP accumulation of meCbl supplementation in wild-type and *tm2345* mutants at D1 stage (mean ± SEM, n=3 biological replicates, animals per condition per replicate = 30, unpaired t-tests: \*\*\**p* < 0.001, ns, no significant differences). Scale bars, 10 μm. Source data are provided as a Source Data file.

*C. elegans*, like humans, cannot synthesize vitamin B12 and thus relies on their bacterial diet. Considering the higher vitamin B12 content in HT115 compared to OP50, and its known role in alleviating cellular stress (particularly oxidative stress)^46,48^, we propose that B12 contributes to the beneficial effects observed in HT115-treated mutants. Indeed, supplementation with 0.15 μM active vitamin B12 (meCbl) significantly reduced *hsp-16.2p::GFP* signals in *lmd-3* mutants (Fig. 6c). Moreover, meCbl treatment during the reproductive period fully restored their fecundity to wild-type levels (Fig. 6d).

Vitamin B12 deficiency impairs canonical propionate metabolism in *C. elegans*, causing propionate accumulation or B12-independent pathway activation (Supplementary Fig. 12c)^49^. Considering that lysosomes are essential for the uptake and processing of vitamin B12 (Supplementary Fig. 12d)^50^, we investigated if *lmd-3* mutants display B12 deficiency-consistent alterations in propionate metabolism. Indeed, high expression of propionate shunt pathway genes, such as *acdh-1* and *hphd-1*, was detected in *tm2345* samples, and this upregulation was rescued by both HT115 and meCbl feeding (Fig. 6e). Furthermore, meCbl supplementation fully restored the mitochondrial membrane potential to wild-type (Supplementary Fig. 12e), counteracting the significant reduction attributed to propionate accumulation^51^. MeCbl’s rescue of propionate metabolism and mitochondrial function in *tm2345* mutants demonstrates that LMD-3 regulates mitochondrial dynamics via vitamin B12 metabolism.

To elucidate the involvement of active vitamin B12 in lysosomal homeostasis, we analyzed the capacity of meCbl supplementation to mitigate lysosomal dysfunction in *lmd-3* mutants. MeCbl supplementation significantly rescued lysosomal acidity defect, as demonstrated by LSG/LTR co-staining (Fig. 6f, Supplementary Fig. 12f) and the pH-sensitive reporter NUC-1::pHTomato (Supplementary Fig. 12g, h). This improvement in lysosomal acidity subsequently led to a significant reduction of the aggregable protein NMY-2::GFP in germline of *tm2345* mutant (Fig. 6g, h), indicating improved lysosomal function, specifically in the context of protein clearance. Together, these results show that vitamin B12 supplementation restores reproductive capacity in *lmd-3* mutants by mitigating cellular stresses and improving lysosomal function.

## Discussion

Our study identifies LMD-3 in *C. elegans*, a conserved LysM domain protein, as a critical regulator of proteostasis and reproductive capacity, offering insights into age-related fertility decline and potential therapeutic avenues. We propose that LMD-3 governs lysosomal acidification via V-ATPase and coordinates with vitellogenin for yolk protein sorting. Additionally, dietary B12, imported through lysosomal degradation of B12-binding proteins, regulates cellular hemostasis and oocyte function (Fig. 7). Collectively, these results define previously unknown physiological functions of LMD-3 in maintaining reproductive health and proteostasis through its interaction with V-ATPase and vitellogenin.

**Fig 7.**
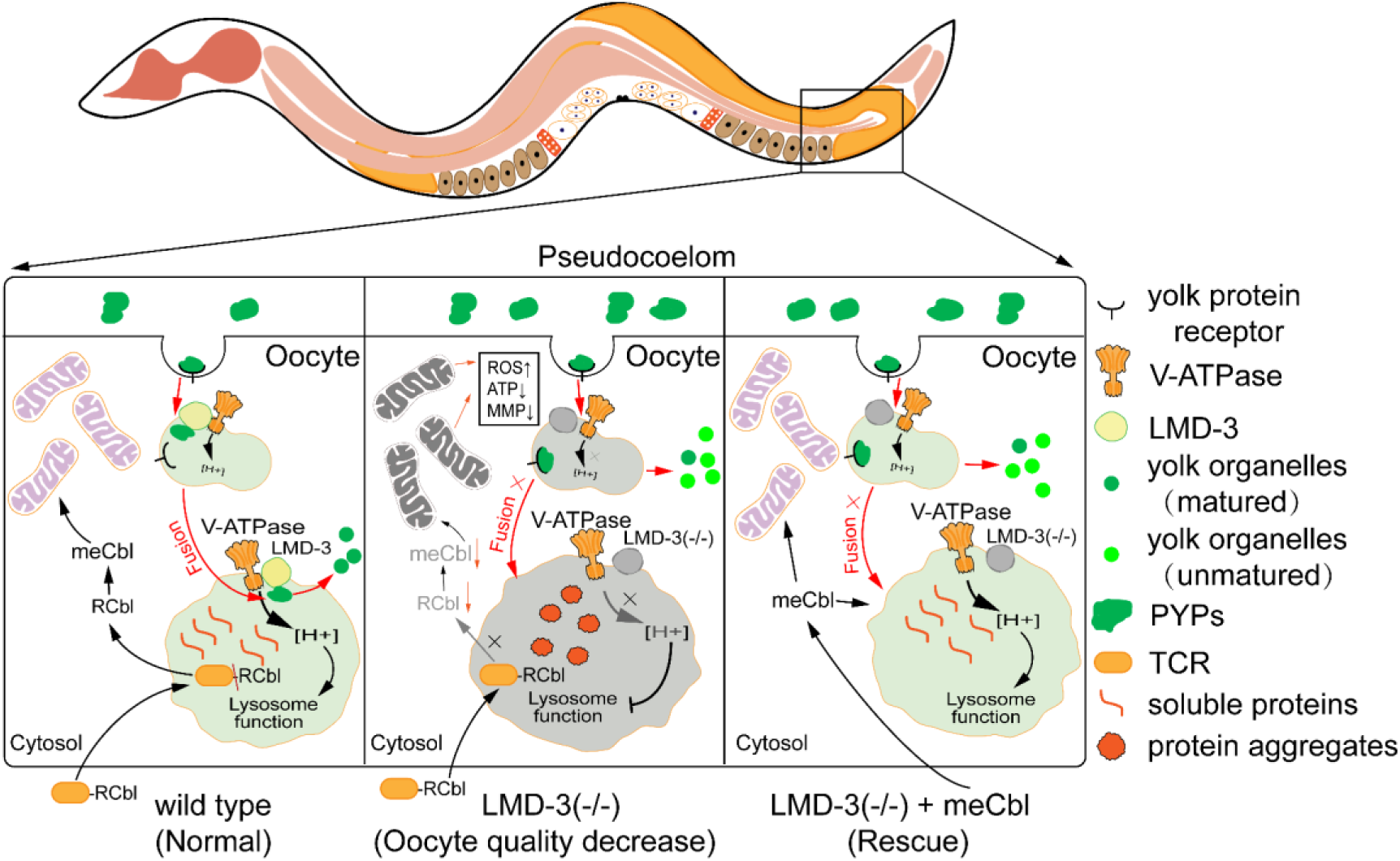
Schematic model showing LMD-3 and vitamin B12 in cellular homeostasis and oocyte function. Under normal conditions, LMD-3 regulates lysosomal acidification (via V-ATPase) and coordinates with vitellogenin for yolk protein sorting, while dietary vitamin B12 is imported via lysosomal degradation of B12-binding proteins and transported to mitochondria and cytoplasm. LMD-3 deficiency increases lysosomal acidification, leading to vitamin B12 deficiency, vitellogenin aggregation, disrupted mitochondrial function, and reduced fertility. Active vitamin B12 (meCbl) supplementation restores reproductive capacity by rescuing the mitochondria-lysosome axis. MMP, mitochondrial membrane potential; PYP, pseudocoelomic yolk patches; TCR, transcobalamin receptor; meCbl, methylcobalamin; RCbl, adenosylcobalamin.

By IP-MS assays, we found that LMD-3 interacts with members of the V-ATPase complex and all vitellogenin proteins. Exhibiting a similar intriguing function, altered expression of *vha-12* (Supplementary Fig. 9a) and other V-ATPase genes mirrored the *lmd-3* loss-of-function phenotypes, including constitutive ER stress response, reduced lysosomal acidity, and defective reproductive capacity in *C. elegans*^12^. While RNAi knockdown of most V-ATPase-related genes (such as *vha-1, vha-2, vha-11, vha-12*, and *vha-13*) caused more severe phenotypes than *lmd-3* deficiency, often leading to lethality or developmental arrest that precluded reproductive analysis, later-stage RNAi treatment in L4 larvae resulted in robust fertility defects (Supplementary Table 3). Notably, *lmd-3* null mutants, despite displaying similar lysosomal and germline defects to these V-ATPase gene knockdowns, remained viable. This suggests that LMD-3 functions in concert with V-ATPase in lysosomal processes but may have more specialized roles, potentially facilitating the trafficking of specific client proteins like vitellogenin.

Consistent with elevated ROS and decreased mitochondrial function observed in POF patients^52,53^, both *lmd-3* loss-of-function and ΔTLDc mutants in *C. elegans* phenocopy these defects. Mechanistically, LMD-3’s conserved TLDc domain is critical for lysosomal acidity and vitellogenin metabolism, as its deletion mirrors *lmd-3* null mutants. Mammalian genomes encode two OXR1 family proteins, OXR1 and NCOA7, both exhibiting oxidation resistance through their conserved TLDc domain. OXR1 is a known susceptibility locus for neurological disorders, primarily due to its roles in lysosomal dysfunction and oxidative stress^17^. The NCOA7b isoform, which also contains a TLDc domain, is highly expressed in reproductive organs, particularly the ovary and uterus^54^. The functional redundancy of OXR1 and NCOA7 in lysosomal acidity or reproductive health in human diseases remains unknown. As *C. elegans* possesses a single *Oxr1* orthologue (Supplementary Fig. 1a), it offers a valuable model for further insights into this protein family’s functions and mechanisms. This divergence suggests evolutionarily distinct proteostasis strategies between nematodes and higher organisms. Future studies on the conserved roles of mammalian OXR1 proteins in health and disease may reveal therapeutic targets.

Our findings align with prior reports that sperm-triggered V-ATPase activation acidifies lysosomes during oocyte maturation, facilitating the clearance of aggregation-prone proteins like NMY-2 and restoring oocyte proteostasis^55^. This mechanism resembles observations in mammals, where lysosome-derived organelles, such as EndoLysosomal Vesicular Assemblies (ELVAs), sustain oocyte proteostasis by eliminating protein aggregates^56^. Importantly, we uncovered that *lmd-3* deficiency drives pathological vitellogenin accumulation in the pseudocoelom, likely through impaired translocation of its receptor RME-2 to the oocyte plasma membrane^57^. This directly impairs lysosomal activity, establishing a causal link between vitellogenin overload and reproductive senescence^34^. Significantly, while autophagy is indispensable for somatic cell and germ cell homeostasis^58^, our data suggest that germline proteostasis in *C. elegans* predominantly relies on lysosomal activity rather than canonical autophagy (data not shown). This specialization may represent an evolutionary optimization to conserve energy in the metabolically demanding reproductive system.

Our study reveals active vitamin B12 (meCbl) as a promising therapeutic strategy for POF, an unmet medical need^59,60^. We found that meCbl supplementation restores oocyte quality and reproductive capacity in *lmd-3* mutants through two synergistic pathways. First, metabolic reprogramming: meCbl, a cofactor for methionine synthase and methylmalonyl-CoA mutase^49,61^, enhances propionyl-CoA metabolism, reducing ROS and restoring mitochondrial membrane potential^62^. Second, lysosomal revitalization: meCbl may directly repair V-ATPase activity^50^, restoring lysosomal acidity and clearing protein aggregates. These effects, mirroring dietary B12 interventions in *C. elegans*, support its use for reproductive decline. However, meCbl couldn’t reverse vitellogenin accumulation in TLDc mutants, indicating LMD-3’s structural integrity is crucial for lysosomal interaction, suggesting a need for combinatorial therapies. Nonetheless, meCbl is a promising POF treatment, especially when lysosomal-mitochondrial coordination is intact. Further studies on the mechanism, biological function, and regulation of Oxr1 family proteins are needed to fully realize these therapeutic potentials.

## Methods

### *C. elegans* culture and strains

*C. elegans* were maintained on standard nematode growth medium (NGM) plates seeded with *E. coli* OP50 at 20°C. Synchronized worm populations were obtained by bleaching gravid adults. The N2 Bristol strain served as the wild-type, and genotypes of all strains used in this study are listed (Supplementary Table 4).

### CRISPR-Cas9 mediated mutagenesis

The precise LysM domain deletion (Δ22-67) and GRAM domain deletion (Δ135-203) of *lmd-3* gene were generated by SunyBiotech (Fuzhou, China). The TLDc domain deletion (Δ627-835) of *lmd-3* gene was generated using established CRISPR-Cas9 methods^63^. PCR and DNA sequencing were used to screen deletion strains (Supplementary Table 1), which were outcrossed to wild-type at least three times before analysis.

### Plasmid construction

The *PCR2.1-nd-1* plasmid was generated by PCR amplifying the *nd-1* fragment from N2 genomic DNA. DNA fragments were inserted into the PCR2.1 vector using Gibson Assembly. The 1XFLAG::LMD-3 expression vectors were generated by PCR amplification of the *lmd-3* cDNA sequence from wild-type and *lmd-3* mutants, followed by subcloning of the amplified DNA fragments into the pcDNA3.1 vector. Primers used for the amplification are listed (Supplementary Table 1).

### Transgenes generation

Transgenic strains in this study were generated as extrachromosomal arrays by standard microinjection method. And UV radiation to generated the integrated line *bmsIs15, bmsIs37,* and *bmsIs48* (Supplementary Table 4).

### Quantitative real-time PCR (qRT-PCR)

For RNA isolation, 200 synchronized L4 stage worms were transferred into Trizol Reagent (Invitrogen, 15596026CN) and snap-frozen in liquid nitrogen. Total RNA was then isolated using standard chloroform extraction and isopropanol precipitation. Subsequently, 1.2 μg of RNA was reverse transcribed by cDNA Synthesis Kit (Vazyme, R312-01). qRT-PCR was performed following the protocol for ChamQ SYBR Color qPCR Master Mix (Vazyme, Q411-02). Transcript levels were normalized to *act-1*.

### UV radiation exposures survival assays

UV radiation exposures were performed using a SCIENTZ03-II UV Crosslinker, which has an emission peak at 365 nm (UVA spectrum). Briefly, stage-synchronized L4 animals were washed and then irradiated on empty agar plates at either 0 or 300 J/m². After irradiation, worms were transferred to NGM plates and cultured at 20°C. Following a 24-hour recovery period on OP50 bacteria, animals were moved to fresh petri dishes every two days until day 12. Worm survival rates were subsequently measured every two days.

### Reproductive capacity analysis

Reproductive capacity was determined by transferring 15 synchronized L4 hermaphrodite larvae to fresh plates every 12 hours until reproduction ceased, a method consistent with previous descriptions^64^. We calculated the total progeny per hermaphrodite. All experiments were performed in triplicate at 20°C. In NH_4_Cl treatment assay, 2M NH_4_Cl was diluted 1:100 into freshly grown OP50 bacteria and spread on NGM plates, after which progenies were counted using the same method.

### Microscopy and Image Analysis

For microscopy, synchronized worms were mounted on 2.5% agar pads containing 5 mM levamisole to facilitate imaging. Images were captured using either a Nexcope 910 or a Carl Zeiss LSM 980 laser scanning confocal microscope. Fluorescence intensity was quantified with ImageJ software.

### Germline staining and quantification

Synchronized D1 adult worms (derived from L4 stage worms transferred to fresh NGM plates 24 hours prior) were fixed in ice-cold methanol for 5 minutes. Following methanol removal, worms were washed three times with M9 buffer (4,000 rpm, 1 minute centrifugation). Samples were then incubated in 500 ng/mL DAPI solution (Macklin, D807022) in the dark for 30 minutes, followed by three additional M9 buffer washes. Worms were mounted for imaging, and their DAPI-stained reproductive gonads were visualized via laser confocal microscopy. ImageJ software facilitated the quantitative analysis of total germ cells and germline stem cells in each gonad.

### SYTO 12 staining for germline apoptotic cells

L4 stage worms from both control and experimental groups were transferred to fresh NGM plates. Synchronized D1 or D2 adults were incubated in the dark with 33 μM SYTO 12 dye (Thermo Fisher Scientific, S7575), which stains germline for 3 hours. After removing the dye, worms were washed by adding 500 μL M9 buffer and centrifuging at 4,000 rpm for 1 minute, followed by discarding the buffer. To clear dye-labeled bacteria from their intestines, worms were transferred to NGM plates for a 1-hour recovery period. Finally, worms were anesthetized with 5 mM levamisole and mounted on 2% agar pads for bright-field and green fluorescence imaging using a 40x objective.

### Dihydroethidium (DHE) staining

Synchronized worms were washed three times with M9 buffer. Subsequently, they were incubated in 100 μL of 10 μM Dihydroethidium (DHE) (Beyotime, S0063) in the dark for 1 hour. After removing the DHE solution, worms were washed three times with 500 μL M9 buffer, centrifuging at 4,000 rpm for 1 minute between washes. The relative DHE fluorescence intensity was quantified using ImageJ software.

### Quantification of ROS content

After three washes with M9 buffer, synchronized worms were lysed by sonication on ice in protein lysis buffer (Beyotime, P0013) containing protease inhibitors (Bimake, B14002) and 1 mM PMSF. Lysates were centrifuged at 15,000 g (4°C for 30 minutes). Thirty microliters of the resulting supernatant were incubated with 30 μL of 100 μM DCFH-DA (Beyotime, S0033S) in the dark for 30 minutes. ROS content per mg of protein was calculated by normalizing the 490 nm absorbance to the protein concentration.

### TMRE fluorescence quantification

TMRE (Beyotime, C2001S) was added to the bacterial lawn from a 10 mM stock solution to a final concentration of 5 mM and dried for 2–3 hours at room temperature. D1 stage animals were then transferred to this prepared bacterial lawn for imaging. To calculate the overall normalized TMRE fluorescence, we determined the ratio of TMRE (Red) to MAI-2::GFP (Green) fluorescence. The wild-type group was normalized to 1.0.

### JC-1 staining and analysis

Isolated mitochondria were resuspended in mitochondrial isolation buffer containing diluted JC-1 staining dye (Beyotime) for 15 minutes. After three washes, mitochondria were transferred to a 96-well black plate. Red fluorescence (Ex: 525 nm, Em: 590 nm) and green fluorescence (Ex: 490 nm, Em: 530 nm) were measured. The mitochondrial membrane potential ΔΨm was calculated as the ratio of red to green fluorescence, normalized to 1.0 for the control group.

### Dual lysosomal staining and quantification

D1 stage worms were incubated in 100 μL of M9 buffer containing 10 mM LysoSensor Green DND 189 (Yeasen, 40767ES50) and LysoTracker Red DND 99 (Yeasen, 40739ES50) for 1 hour at 20°C in the dark to stain lysosomes. Following staining, worms were transferred to NGM plates with fresh OP50 and allowed to recover for 1 hour at 20°C in the dark before examination. The relative intensity ratio of LSG to LTR was quantified using ImageJ software.

### Cell culture and Transient transfection

HEK-293T cells were cultured in Dulbecco’s Modified Eagle’s Medium (DMEM) (Gibco, C119955-00BT) supplemented with 10% (v/v) Fetal Bovine Serum (FBS) (Excell, FSP-500). Cells were maintained at 37°C in a humidified incubator with 5% CO2. For transfection, 1 µg of total DNA per 10^6 cells was used with Polyethyleneimine (PEI) (Sigma-Aldrich, 408727) at a DNA:PEI ratio of 1:4.5.

### Cycloheximide pulse chase assay

We assessed the protein stability of 1xFLAG-tagged LMD-3 deletion constructs in HEK293T cells using a CHX (cycloheximide) pulse-chase assay, as previously described^65^. Thirty-six hours post-transfection, cells were treated with either 50 µg/mL CHX for the indicated time points. We then collected the cells and analyzed protein levels via Western blotting.

### Quantification of mtDNA copy number and lesions

Synchronized worms were analyzed for mitochondrial DNA (mtDNA) copy number and damage. Six worms were lysed in 90 μL of lysis buffer by freezing at -80°C for 1 hour, then heating at 65°C for 1 hour and 95°C for 15 minutes^66^. Long amplicon PCR was subsequently performed. mtDNA copy number was quantified using a plasmid-based standard curve and real-time PCR, normalized to the six individuals per sample due to the consistent somatic cell number in worms. mtDNA lesions were calculated from long-amplicon PCR products, normalized to mtDNA copy number and relative to the control within each strain. Each biological sample was amplified in triplicate, and the experiment was repeated twice.

### RNA interference (RNAi) protocol

The RNA interference (RNAi) experiments utilized an RNA interference competent OP50 (iOP50) *E. coli* strain, which was transformed with either the L4440 RNAi control vector or the *vit-2* RNAi vector. For RNAi treatment, iOP50 expressing the desired vector was grown overnight at 37°C in LB broth supplemented with 100 µg/mL ampicillin, which was then spread onto NGM plates containing 100 µg/mL ampicillin and 1 mM isopropyl β-D-1-thiogalactopyranoside (IPTG) to induce dsRNA expression. The seeded plates were incubated overnight at 25°C to allow bacterial growth and dsRNA production. Afterward, synchronized L4 *C. elegans* larvae were carefully moved to the RNAi-seeded plates. The worms were then cultured at 20°C, until they reached their young adult stage. Young adult worms were then collected for downstream experiments.

### Mecobalamin supplementation

A 150 µM stock solution of meCbl (Macklin, M812748) was prepared in DMSO. For experiments, this stock was diluted to a final concentration of 0.15 µM in NGM agar.

### Quantification of NUC-1::pHTomato intensity

*C. elegans* D1 adults expressing P*_hs_*NUC-1::pHTomato were incubated at 33°C for 30 minutes, then allowed to recover at 20°C for 24 hours before examination. The average pHTomato fluorescence intensity per hypodermal area was quantified using ImageJ software. At least 20 worms were analyzed per strain.

### Quantification of NUC-1::sfGFP::mCherry number

*C. elegans* D1 adults expressing P*hsp*NUC-1::sfGFP::mCherry was incubated at 33°C for 30 min, and then the worms were captured by laser scanning confocal microscopy after 2, 12 and 24 hours. The number of NUC-1::mCherry puncta per area in the hypodermis was measured by Image J software.

### Mitochondrial isolation

We manually collected approximately 10,000 animals and washed them three times with M9 buffer. Worms were then resuspended in mitochondrial isolation buffer (200 mM Mannitol, 50 mM Sucrose, 10 mM KCl, 1 mM EDTA, 10 mM HEPES-KOH (pH 7.4), 0.1% BSA, and protease inhibitors (Bimake, B14002). Homogenization occurred on ice in a 5 mL Dounce homogenizer with 20 gentle strokes. The homogenate was first centrifuged at 1,000 g for 10 minutes to remove debris. The resulting supernatant was then centrifuged at 12,000 g for 10 minutes to obtain a crude mitochondrial pellet, from which the supernatant was carefully decanted.

### Quantification of ATP levels

The ATP content of worms and isolated mitochondria was determined using the ATP-dependent Firefly Luciferase Bioluminescence Assay Kit (Beyotime, S0026). For whole worm samples, 200 worms were collected, washed into M9, and lysed via sonication on ice in 100 µL lysis buffer, followed by centrifugation (12,000 g, 5 min, 4°C). Twenty microliters of the resulting supernatant, or isolated mitochondria resuspended in ATP reagent, were then subjected to the luciferin-luciferase assay according to the kit’s protocol. Luminescence was read on a Centro XS^3^ LB 960 (Berthold). ATP content for each sample was derived from a standard curve. Each sample type included three biological replicates.

### Immunoprecipitation and Protein Identification

Worms were washed thrice with M9 buffer before lysis by sonication on ice in protein lysis buffer (Beyotime, P0013) supplemented with protease inhibitors (Bimake, B14002) and 1 mM PMSF. Lysates were then centrifuged at 15,000 g for 30 minutes at 4°C. The supernatant was pre-cleared with binding control magnetic agarose beads (ChromoTek, bmab-20) for 20 minutes at 4°C. Subsequently, RFP-Trap magnetic agarose beads (ChromoTek, rtma-10) were incubated with the pre-cleared supernatant for 2 hours at 4°C on a rotating shaker. Beads were washed thrice with PBS, and proteins were eluted by resuspending in 1x sample buffer and heating at 95°C for 10 minutes. Eluted proteins were separated by SDS-PAGE, and gels were silver stained (Beyotime, P0017S). Finally, bands of interest were identified via mass spectrometry.

### Co-immunoprecipitation (Co-IP)

For co-immunoprecipitation, 1 mL of supernatant was first pre-cleared with binding control magnetic agarose beads for 20 minutes at 4°C on a rotating shaker. The pre-cleared supernatant was then incubated for 1 hour at 4°C on a rotating shaker with either GFP-Trap (ChromoTek, rtma-10) or FLAG (Thermo, A36797) magnetic agarose beads. Beads were washed three times with 1X PBS buffer, and proteins were eluted by resuspending in 1x sample buffer and heating at 95°C for 10 minutes. Eluted protein lysates were then analyzed by Western blotting using anti-mCherry rabbit (abcam, ab213511) and anti-GFP rabbit pAb (abclonal, AE011) antibodies.

### Statistical analysis

All data were analyzed using GraphPad Prism 9. The Y-axis error bars represent the Standard Error of the Mean (SEM). We used Student’s t-tests to compare the means of different samples. Statistical significance was defined as *p* < 0.05 and indicated by a single asterisk (*). Double asterisks (**) denoted *p* < 0.01, and triple asterisks (***) indicated *p* < 0.001.

### Data availability

All data in the main Manuscript and Supplementary information are listed in the Source data file. All reagents and strains generated by this study are available through a request to the corresponding author with a completed Material Transfer Agreement. Source data are provided in this paper.

## Acknowledgements

We thank the Caenorhabditis Genetics Center (CGC), Dr. Xiaochen Wang (Southern University of Science and Technology), and Dr. Chenggang Zou (Yunnan University) for strains; This study was supported by National Natural Science Foundation of China (32170781 to Z.Z.), and Natural Science Foundation of Shandong Province (2023HWYQ-014, tsqn202306056, ZR2021QC023, and 2021JK032 to Z.Z.).

## Author contributions

Y.Z., T.W., W.L. and Z.Z. conceived the project, designed and performed the experiments, and analyzed the data. J.H. constructed the plasmids, and conducted oxidative stress assays. M.W. carried out the reproductive assays and analysis of the mutants. M.G. performed the co-immunoprecipitation (co-IP) assays for LMD-3 and VIT-2. Y.Z. and T.W. wrote the manuscript. W.L. and Z.Z. contributed to manuscript editing.

## Competing interests

The authors declare no competing interests.

**Correspondence and requests for materials** should be addressed to Zhe Zhang.

