## Supplementary Figure1 for "Lysosomal control of proteostasis and reproductive capacity by conserved LMD-3 protein in *C. elegans*"

**Supplementary Figures. 1 to 12**

**Supplementary Fig. 1**

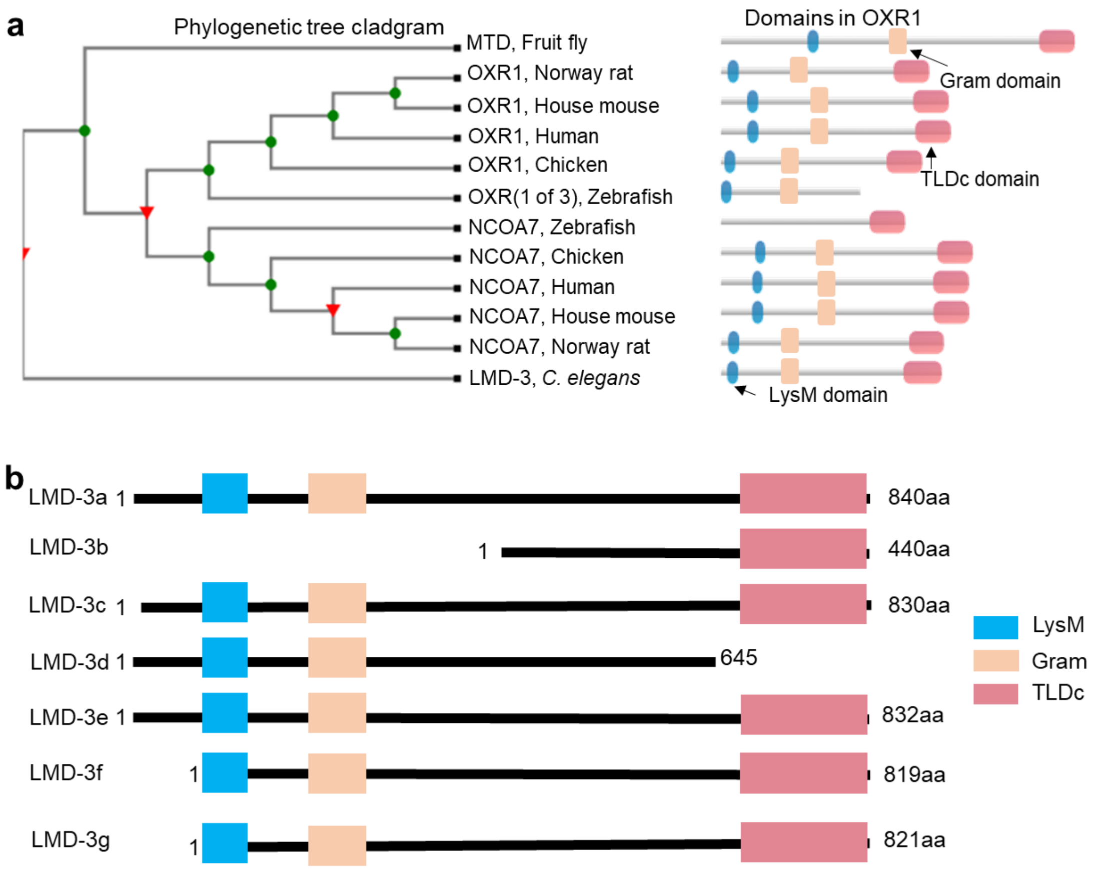


**Supplementary Fig. 1 Evolutionary conservation of OXR1 and *lmd-3* isoform domains. a** Cladogram of phylogenetic tree and conserved domains for the OXR1 protein family from major metazoan species (adapted from www.treefam.org). **b** Predicted peptides and are shown for all *lmd-3* isoforms. The number of amino acids (aa) for each isoform is indicated.

**Supplementary Fig. 2**

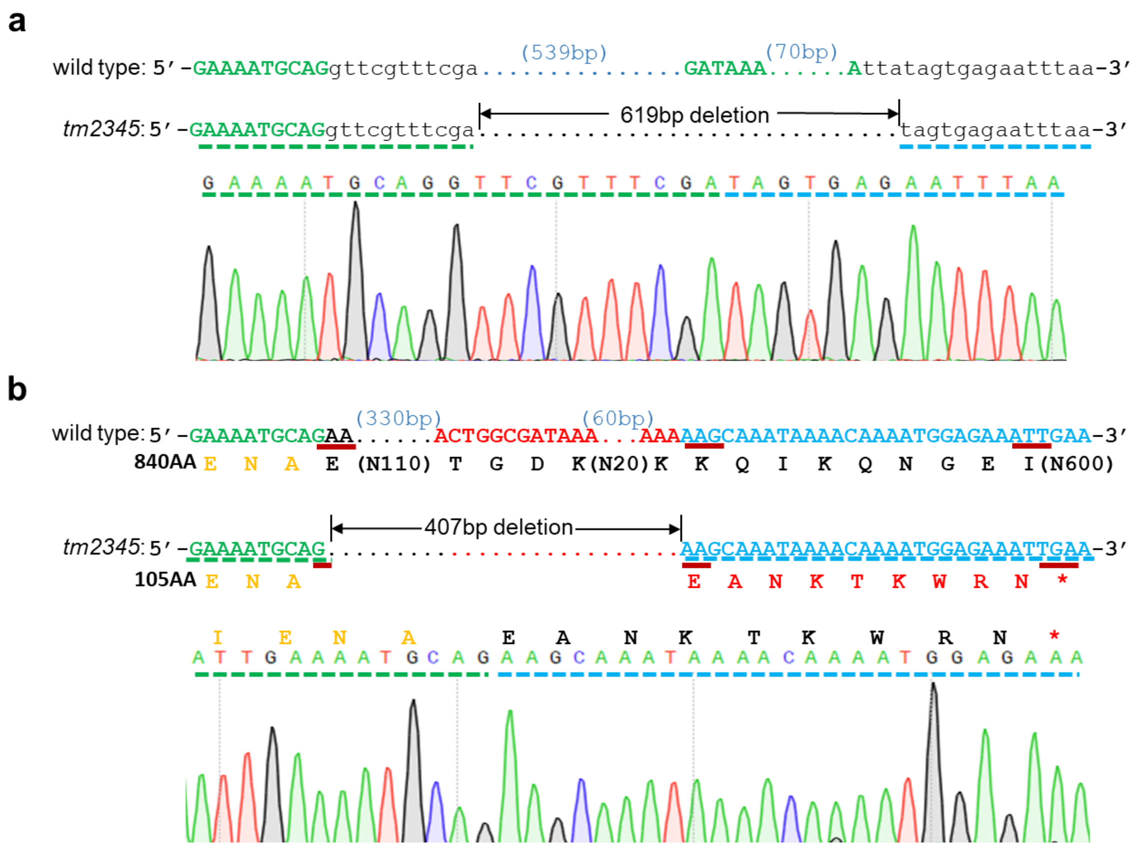


**Supplementary Fig. 2 Sequencing result of *lmd-3(tm2345)*.** The sequence of wild type animals*, lmd-3(tm2345)* animals, and sequencing tracks of *lmd-3(tm2345)* animals near the *tm2345* deletion sites are shown with genomic DNA sequence (**a**) and cDNA sequence (**b**), separately. Uppercase letters indicate coding sequences, while lowercase letters represent introns. Residues with altered amino acid sequences are labeled in red, and the location of the premature stop codon is indicated by a red asterisk.

**Supplementary Fig. 3**

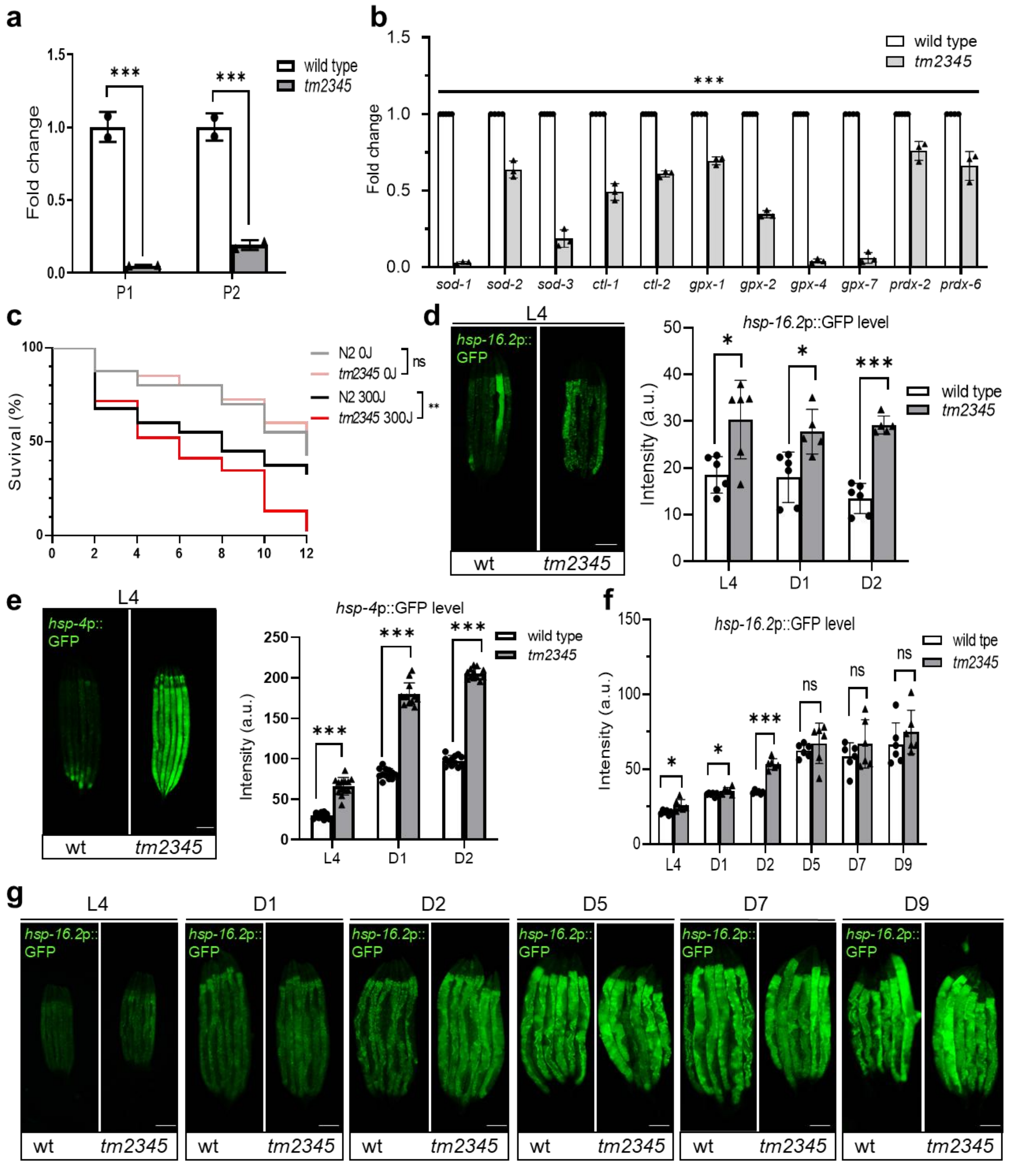


**Supplementary Fig. 3 LMD-3 deficiency induces oxidative stress in *C. elegans*. a, b** Independent repeats of qRT-PCR measurements of mRNA fold changes in wild-type and *tm2345* mutants for endogenous *lmd-3* (n = 2 for each group, unpaired t-tests: ***$p < 0.001$) (**a**) and eleven selected antioxidant genes (n = 3 for each group, unpaired t-tests: ***$p <$

0.001) (**b**). **c** Independent repeat of survival curves of wild-type and *tm2345* animals (n = 30 for each group, unpaired t-tests: **P < 0.01, ns, no significant differences) after UV irradiation (300J/m$^2$). **d, e** Exemplar fluorescence images and quantification for *hsp-16.2*p::GFP (**d**) and *hsp-4*p::GFP (**e**) with wild-type and *tm2345* mutants at indicated stage (n ≥ 6 for each group, unpaired t-tests: *$p < 0.05$, ***$p < 0.001$). L4, the fourth and final larval stage. D1, Day 1 of adulthood. D2, Day 2 of adulthood. **f, g** Cytoplasmic stress reporter *hsp-16.2*p::GFP in wild-type and *tm2345* mutants (n = 6 for each group, unpaired t-tests: *$p < 0.05$, ***$p < 0.001$, ns, no significant differences) throughout the *C. elegans* life cycle with exemplar images (**f**) and quantification (**g**). a.u., arbitrary units. Scale bars represent 100 μm. Source data are provided as a Source Data file.

**Supplementary Fig. 4**

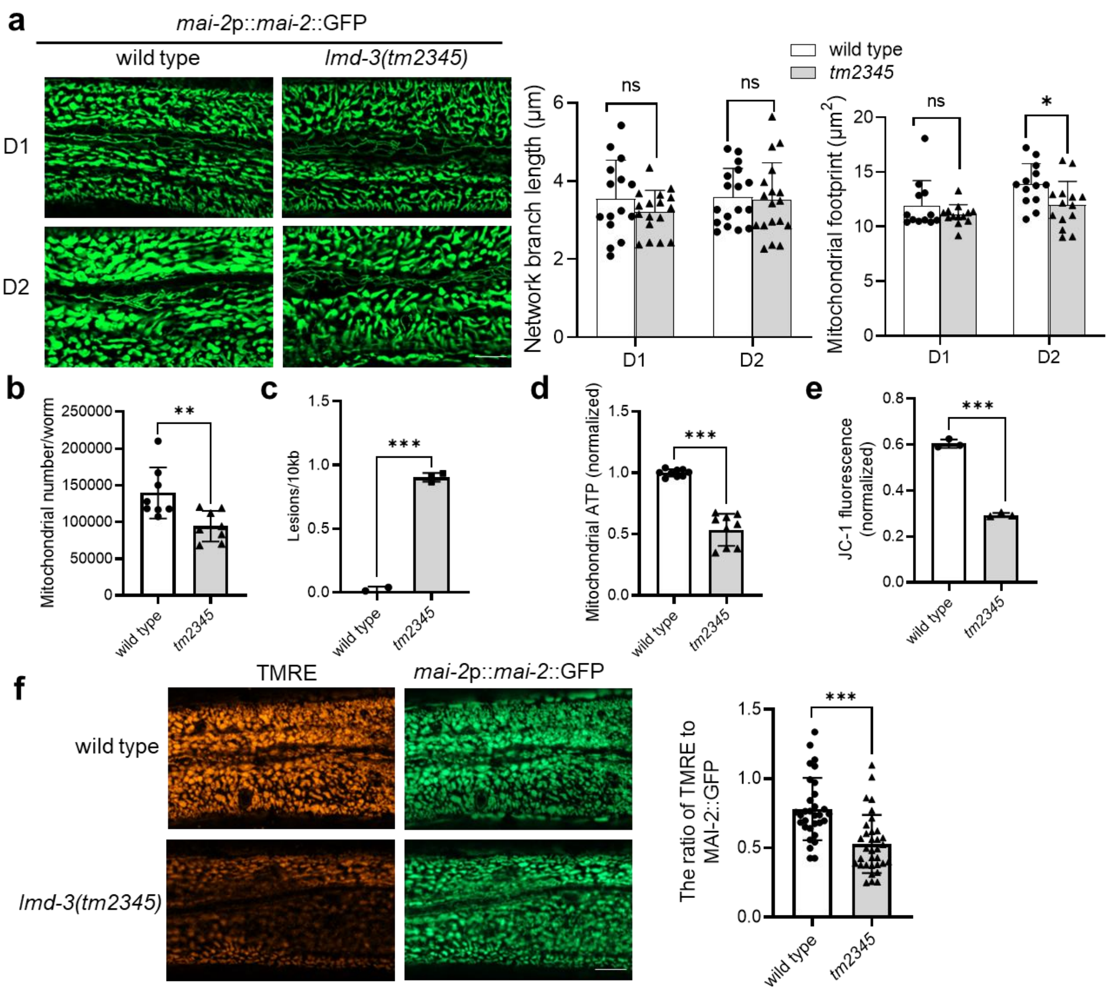


**Supplementary Fig. 4 *lmd-3* deficiency leads to mitochondrial dysfunction in D1 young *C. elegans*. a** Mitochondrial morphological change with *mai-2*p::*mai-2*::GFP reporter in wild-type and *tm2345* animals with Confocal image and quantifiable measurements (n ≥ 12 for each group, unpaired t-tests: *$p$ < 0.05, ns, no significant differences), including network branch length (μm) and mitochondrial footprint (μm$^2$). Scale bars,10 μm. **b** Mean mtDNA copy number in wild-type and *tm2345* mutants (n ≥ 16 for each group, unpaired t-tests: **$p$ < 0.01). **c** Quantification of DNA damage in mitochondria of wild-type and *tm2345* mutants (n = 3 biological replicates, unpaired t-tests: ***$p$ < 0.001). **d** ATP level assessment in crude isolated mitochondria of wild-type and *tm2345* mutants (n = 9 for each group, unpaired t-tests: ***P < 0.001). **e** Quantitative analysis of mitochondrial membrane potential (ΔΨm) in isolated mitochondria from wild-type and *tm2345* mutants (n = 3 biological replicates, unpaired t-tests: ****p* < 0.001). **f** Confocal image and quantification of mitochondrial membrane potential in wild type and *tm2345* mutant nematodes (n ≥ 15 for each group, unpaired t-tests: ****p* < 0.001). Scale bars,10 μm. Source data are provided as a Source Data file.

**Supplementary Fig. 5**

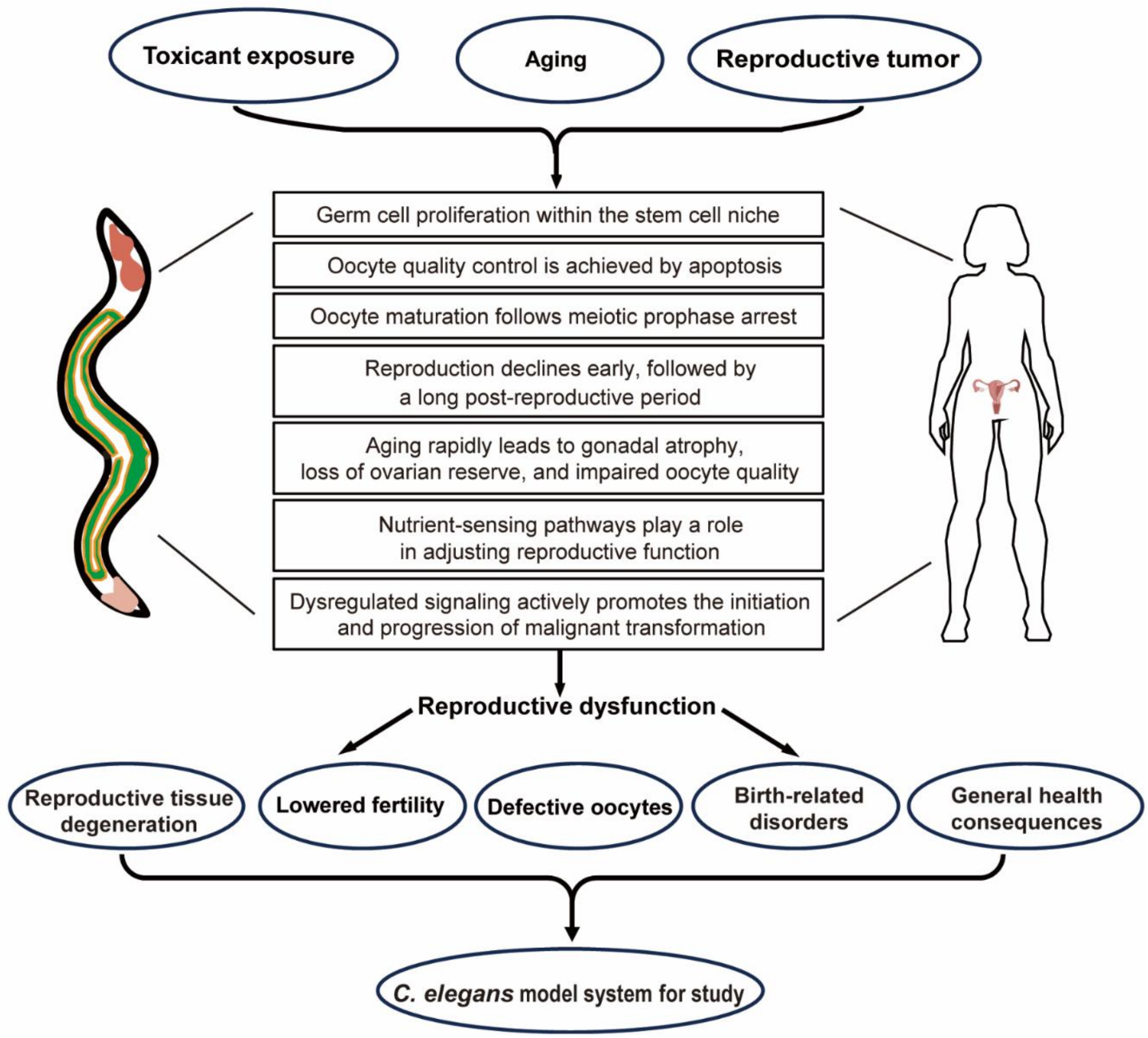


**Supplementary Fig. 5** *C. elegans* as a model for conserved reproductive mechanisms. In both *C. elegans* and humans, toxicant exposure, aging, and reproductive tumor can impair the reproductive system, leading to reproductive dysfunction. Stem cell niches support germline proliferation, while apoptosis ensures oocyte quality. Oocyte maturation follows meiotic prophase arrest. Early reproductive decline precedes a long post-reproductive phase. Aging accelerates gonadal atrophy, ovarian reserve depletion, and oocyte damage. Nutrient sensing modulates reproductive function, and dysregulated signaling can cause malignant transformation. These factors lead to reproductive dysfunction (e.g., reproductive tissue degeneration, reduced fertility, abnormal oocytes, birth defects, and systemic health effects), which is readily studied in *C. elegans*.

**Supplementary Fig. 6**

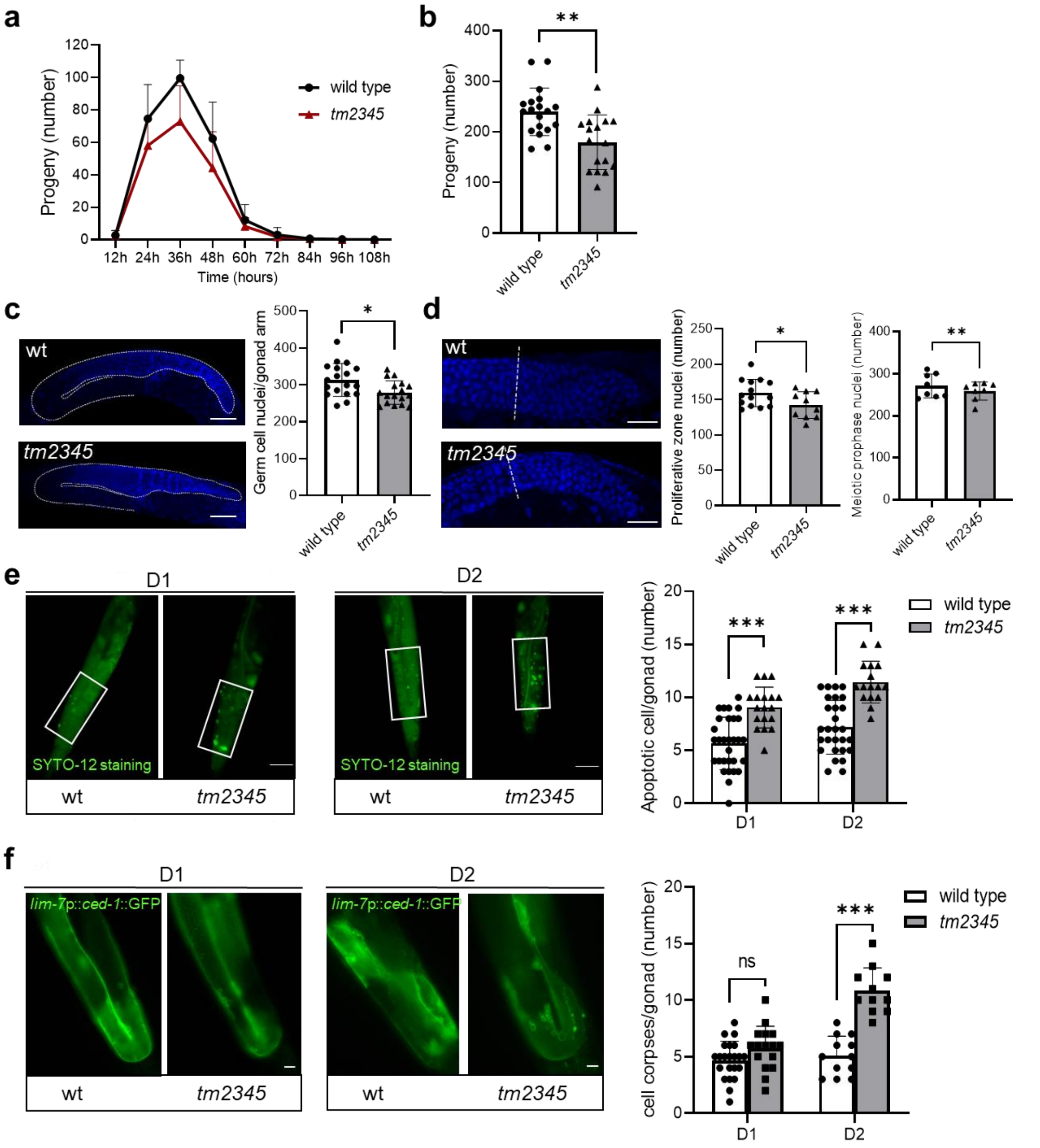


**Supplementary Fig. 6 *lmd-3* deficiency causes germ cell apoptosis and reduced fecundity in *C. elegans*. a** The Independent repeat of progeny production curves of wild-type (black) versus *tm2345* (red) hermaphrodites that survive beyond their reproductive span. **b** Independent repeat of total brood sizes indicates the ability of *C. elegans* to produce offspring in wild-type and *tm2345* (n ≥ 17 for each group, unpaired t-tests: $**p <$

0.01). **c** Independent repeat of exemplar fluorescence images and quantification of DAPI staining for total germline cells with wild-type and *tm2345* mutants (n = 17 for each group, unpaired t-tests: $*p < 0.05$). Scale bars, 50 μm. **d** Independent repeat of exemplar fluorescence images and quantification of DAPI staining for germline stem cells and meiotic cells in wild-type and *tm2345* mutants (n ≥ 8 for each group, unpaired t-tests: $*p < 0.05$, $**p < 0.01$). Scale bars, 20 μm. **e, f** Independent repeat of exemplar fluorescence images and quantification for SYTO12 staining (**e**) and *lim-7*p::*ced-1*::GFP (**f**) with wild-type and *tm2345* mutants at indicated stage (n ≥ 5 for each group, unpaired t-tests: $*p < 0.05$, $**p < 0.01$, $***p < 0.001$). D1, Day 1 of adulthood. D2, Day 2 of adulthood. Scale bars, 50 μm. Source data are provided as a Source Data file.

**Supplementary Fig. 7**

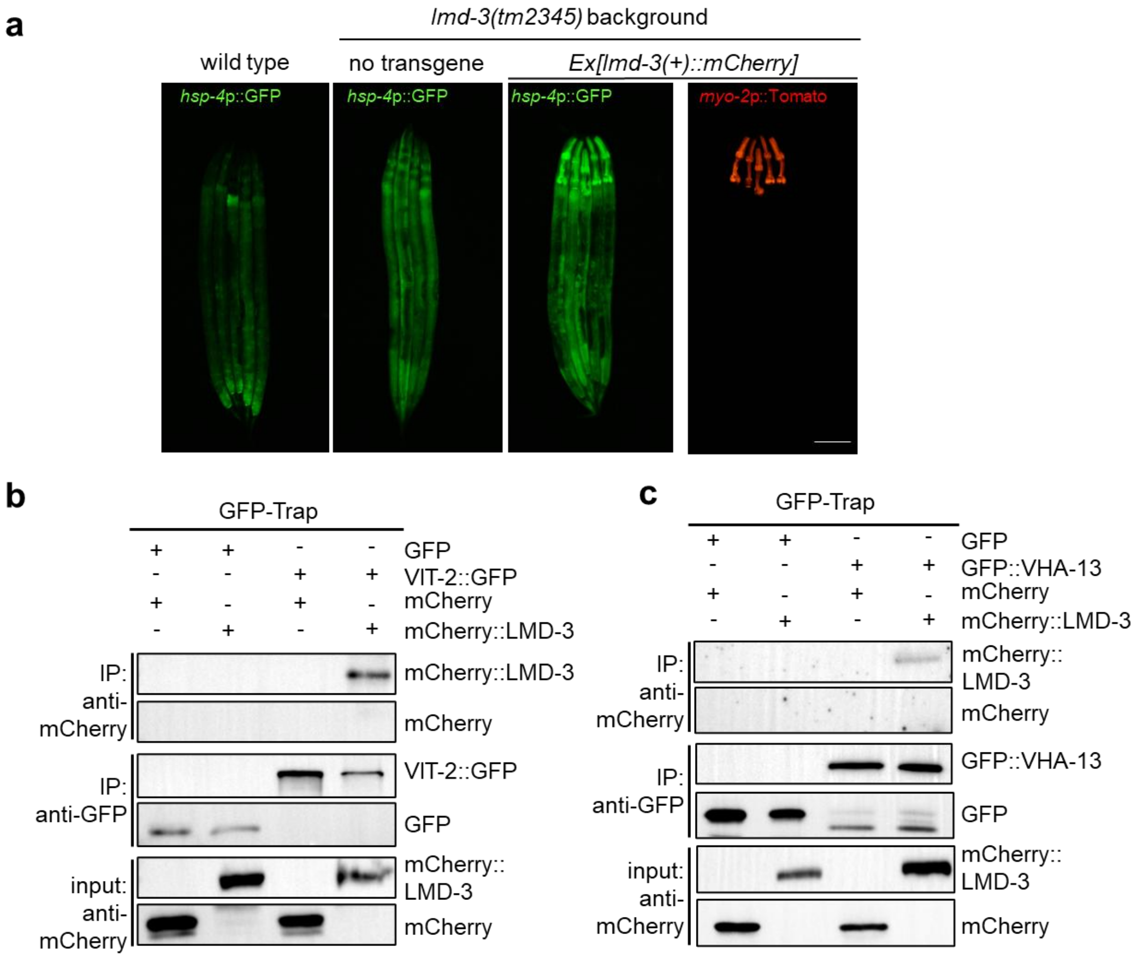


**Supplementary Fig. 7 The interaction of LMD-3 with vitellogenin and V-ATPases is implicated in cellular homeostasis regulation. a** Representative fluorescence images showing that ubiquitously expressed *rpl-28p::lmd-3::mCherry* failed to rescue *hsp-4p*::GFP induction in *lmd-3(tm2345)* mutants. The transgene is marked by pharyngeal *myo-2*p::mCherry. Scale bars, 50 μm. **b, c** Co-Immunoprecipitation (co-IP) and western blot showing biochemical interaction of N-terminal mCherry-labeled LMD-3 with GFP-labeled VIT-2 (**b**) and GFP-labeled VHA-13 (**c**), respectively. Transgenic worms expressed with expression vectors, lysed for immunoprecipitation by GFP-TRAP, and blotted by antibodies against GFP and mCherry.

**Supplementary Fig. 8**

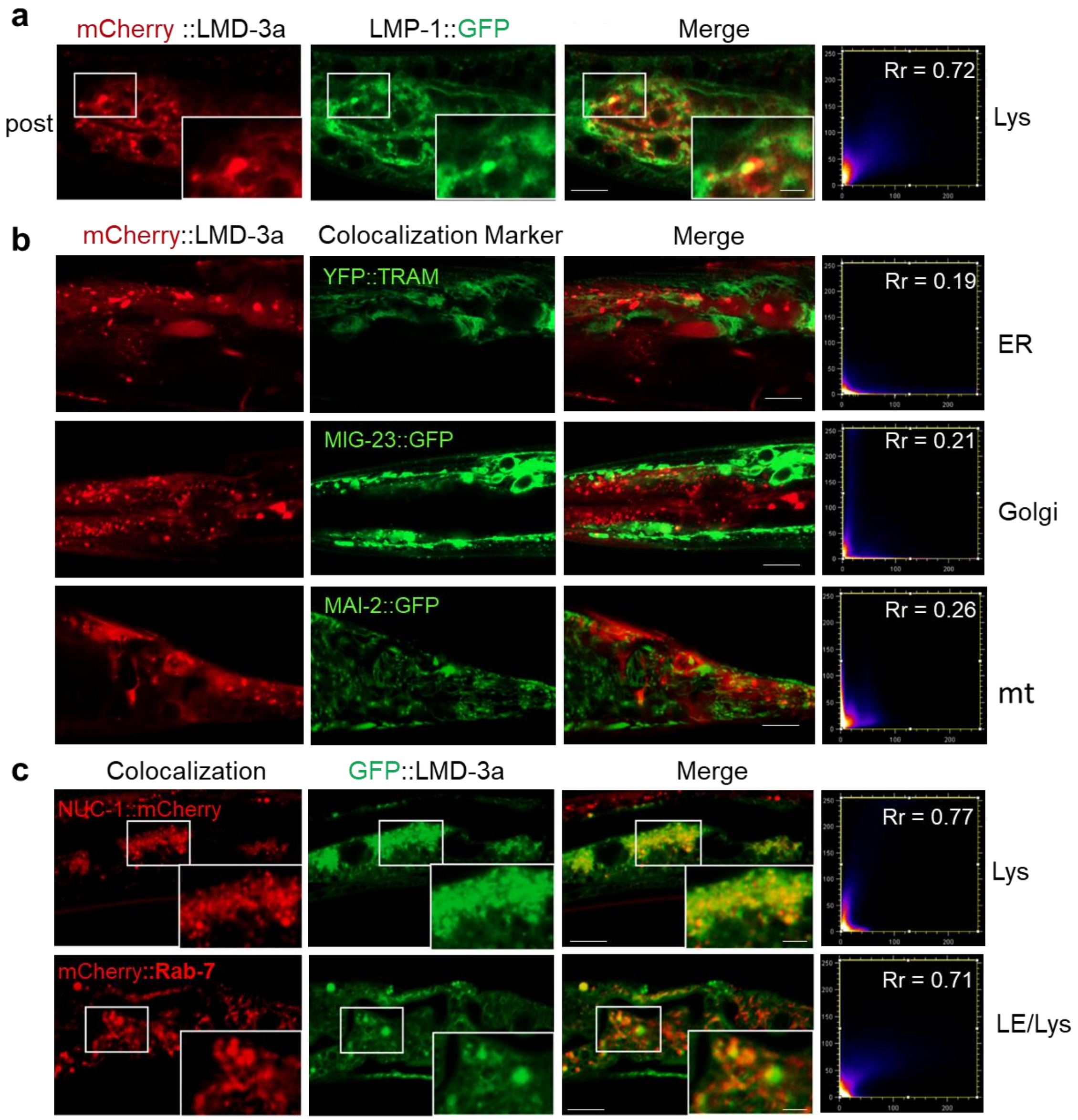


**Supplementary Fig. 8** Subcellular localization of LMD-3 with different fluorescent makers. **a, b** Confocal images showing colocalization of mCherry::LMD-3 with the lysosomal marker in the posterior (post) region (**a**), and markers for the endoplasmic reticulum (ER), golgi apparatus, and mitochondria (mt) in the *C. elegans* hermaphrodite. **c** Confocal images showing colocalization of GFP::LMD-3 with NUC-1::mCherry (lysosomal marker), and late endosome/lysosome maker mCherry::Rab-7. Pearson's correlation coefficient (Rr)

indicates the strength and direction of a linear relationship in pixel intensities (Rr > 0.5 suggests good colocalization). Scale bar, 10 μm. Lys, lysosome. LE, late endosome.

**Supplementary Fig. 9**

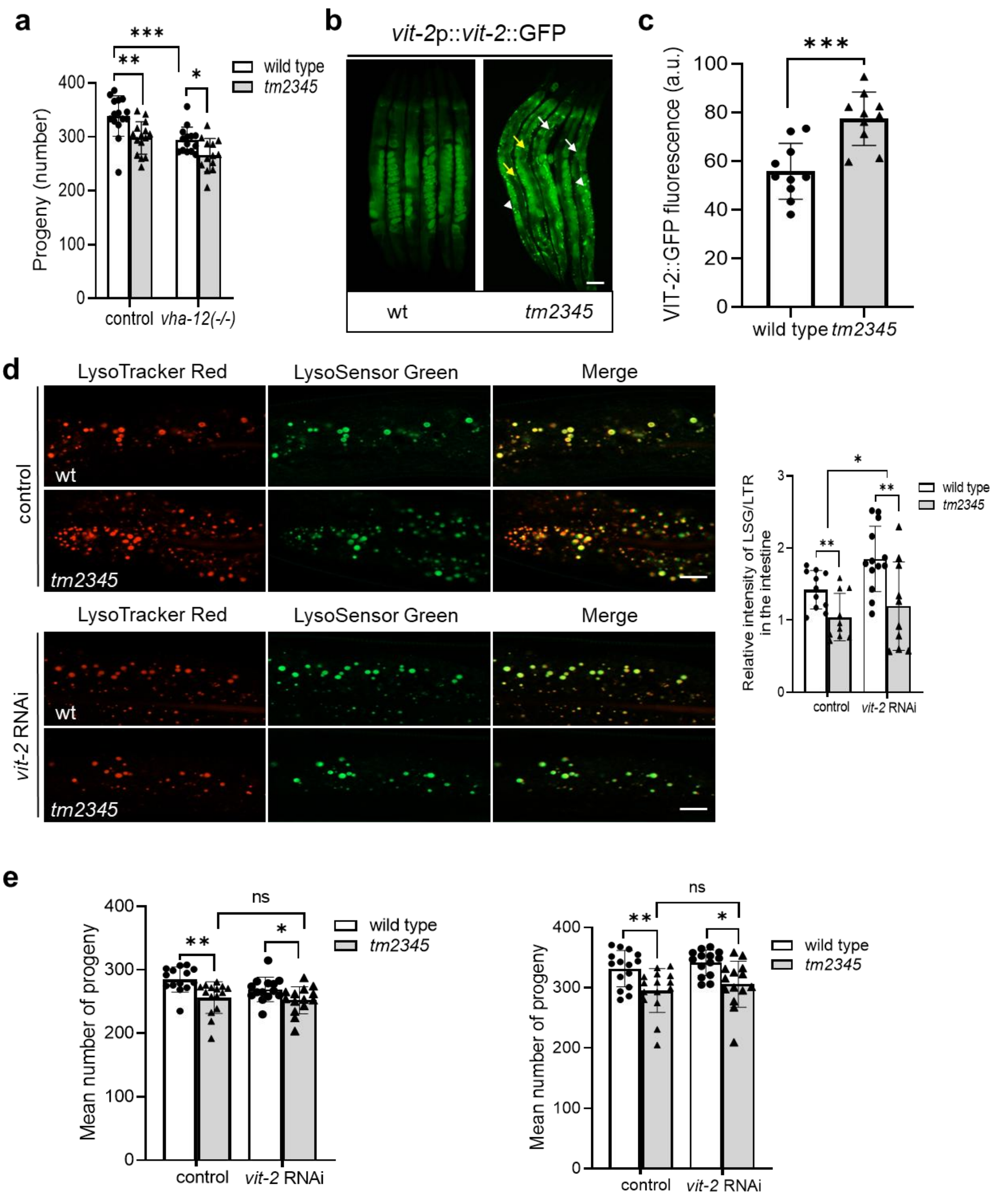


**Supplementary Fig. 9** ***vha-12*** **loss and vitellogenin accumulation synergistically impair reproduction. a** Total brood sizes indicate the effect of *vha-12(ok821)* mutantation on reproductive capacity in wild-type and *tm2345* mutants (n ≥ 13 per group, unpaired t-tests: *$p < 0.05$, **$p < 0.01$, ***$p < 0.001$). **b, c** Exemplar fluorescence images (**b**) and quantification (**c**) *vit-2*p::*vit-2*::GFP accumulation in wild-type and tm2345 mutants at D1 stage (n ≥ 10 for each group, unpaired t-tests: ***$p < 0.001$). Aggregates in different tissues: white arrowheads denote germline, white arrows denote pseudocoelom, yellow arrows denote intestines. a.u., arbitrary units. Scale bars, 50 μm. **d** Confocal fluorescence images (**d**) and quantification (**e**) of LSG DND-189/LTR DND-99 co-stained intestines of control and *vit-2* RNAi-treated wild type and *lmd-3(tm2345)* animals at the D1 stage (n ≥ 10 per group, unpaired t-tests: *$p < 0.05$, **$p < 0.01$). Scale bars, 50 μm. **e** Independent repeats of reproductive capacity of wild-type and *lmd-3(tm2345)* animals with control and *vit-2* RNAi treatment (n ≥ 15 per group, unpaired t-tests: *$p < 0.05$, **$p < 0.01$, ns, no significant differences). Source data are provided as a Source Data file.

**Supplementary Fig. 10**

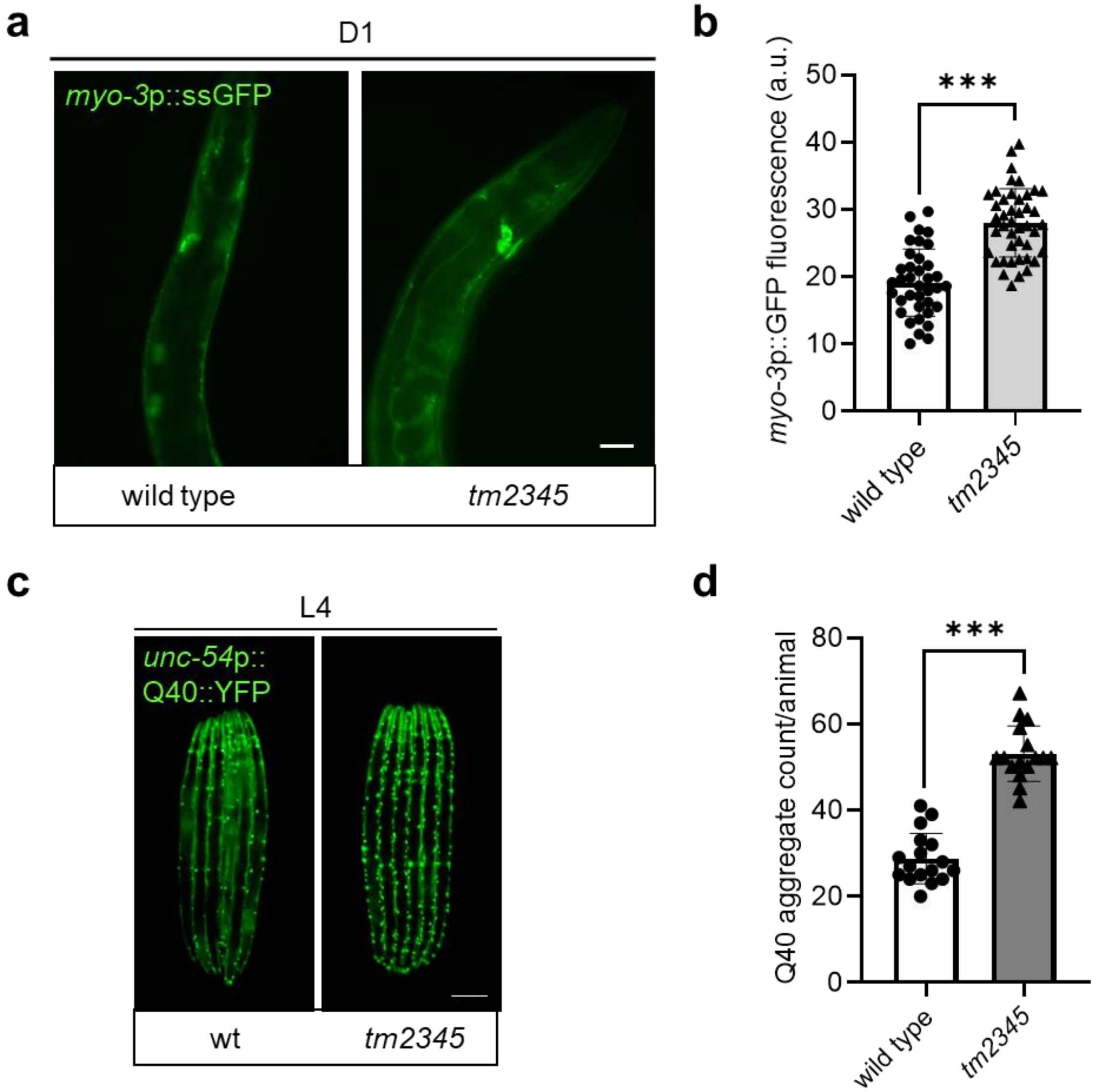


**Supplementary Fig. 10 LMD-3 deficiency impairs lysosomal degradation and proteostasis. a, b** Exemplar fluorescence images (**a**) and quantification (**b**) *myo-3*p::ssGFP accumulation in coelomocytes of wild-type and *tm2345* animals at D1 stage (n ≥ 37 for each group, unpaired t-tests: ***$p < 0.001$). Scale bars, 50 μm. **c, d** Exemplar fluorescence images (**c**) and quantification (**d**) of Q40::YFP aggregation with wild-type and *tm2345* animals at L4 stage (n ≥ 16 for each group, unpaired t-tests: ***$p < 0.001$). Scale bars, 50 μm. a.u., arbitrary units. Source data are provided as a Source Data file.

**Supplementary Fig. 11**

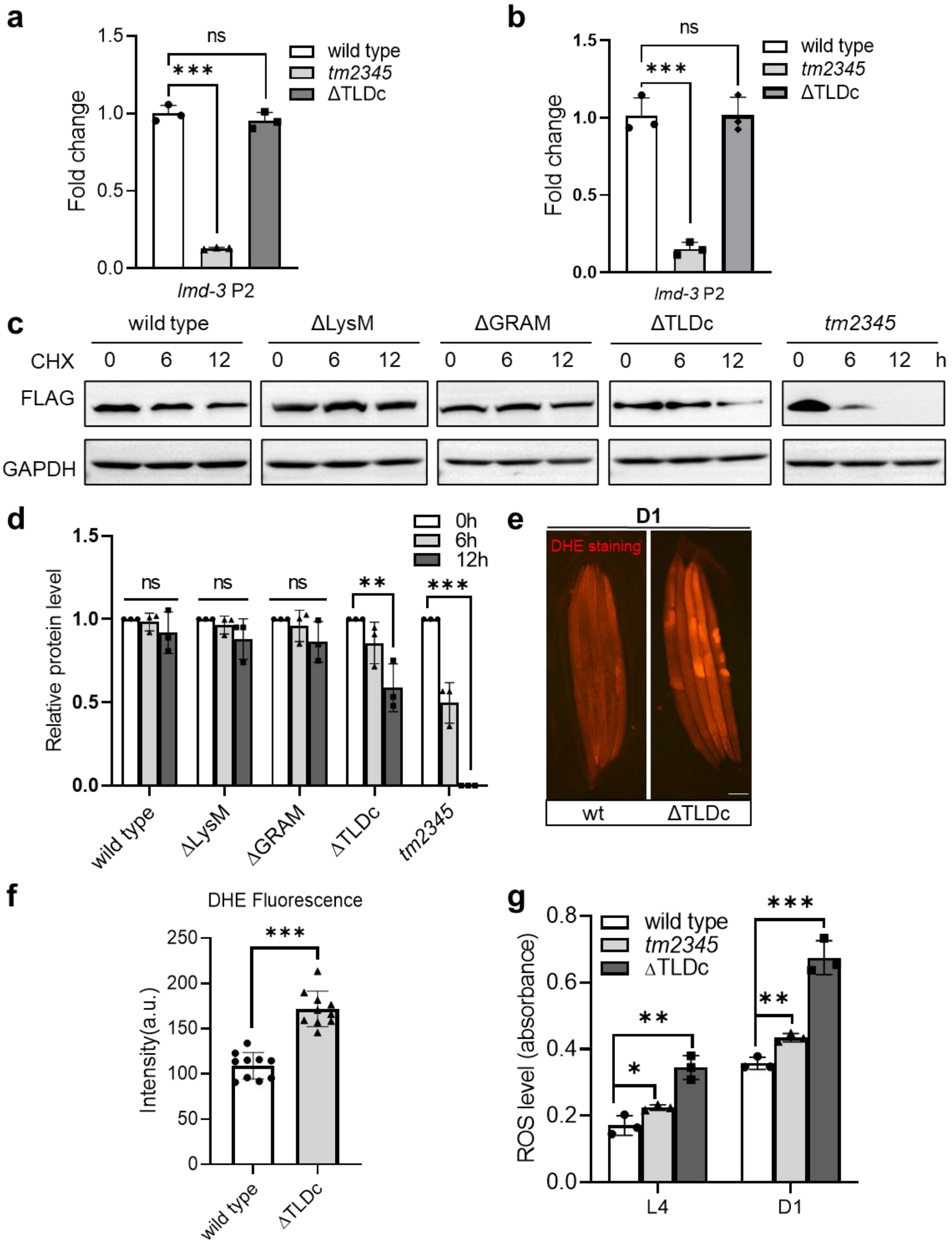

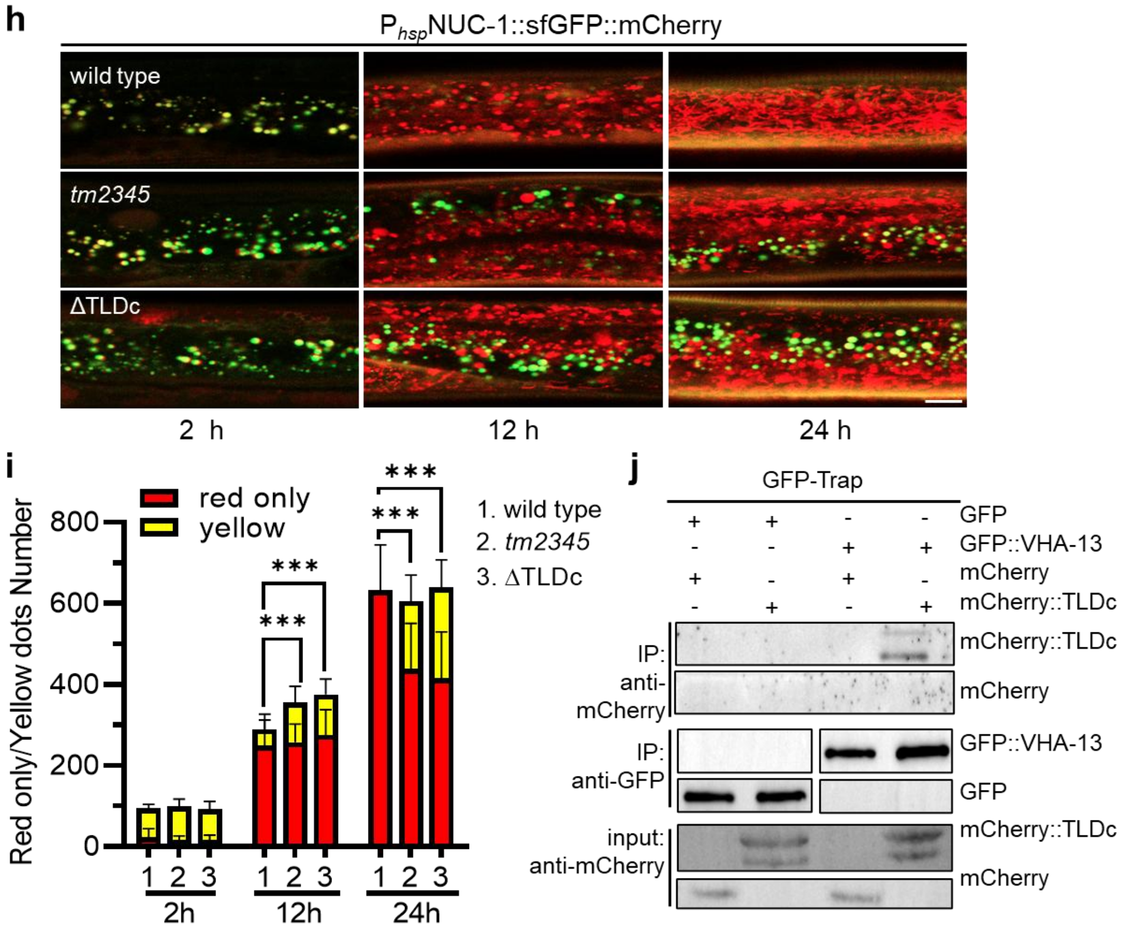


**Supplementary Fig. 11 TLDc domain are essential for LMD-3 functions. a, b** qRT-PCR measurements of mRNA fold changes in wild-type, *tm2345* and ΔTLDc mutants (n = 3 for each group, unpaired t-tests: ***$p$ < 0.001, ns, no significant differences) for endogenous *lmd-3* (**a**) and independent repeats (**b**). **c, d** 1xFLAG-tagged LMD-3 protein turnover with exemplar western blot images (**c**) and semi-quantitative analysis by scanning of blot scanning (**d**) of H293T cells treated with 50 μg/ml cycloheximide (CHX) for 0, 6, and 12 h (n = 3 biological replicates, unpaired t-tests: **$p$ < 0.01, ***$p$ < 0.001, ns, no significant differences). a.u., arbitrary units. **e, f** Superoxide anion ($O_2^-$) level detection by DHE staining with exemplar images (**e**) and quantification (**f** of DHE fluorescence in wild-type and ΔTLDc (n = 10 for each group, unpaired t-tests: ***$p$ < 0.001). Scale bars, 100 μm. a.u., arbitrary units. **g** Reactive oxygen species (ROS) levels detected using

Dichloro-dihydro-fluorescein diacetate (DCFH-DA) assay in different groups (n = 3 biological replicates, unpaired t-tests: *$p < 0.05$, **$p < 0.01$, ***$p < 0.001$). **h, i** Confocal fluorescence images (**h**) and quantification (**i**) the of red-only and yellow puncta of NUC-1::sfGFP::mCherry at various time points after heat shock in wild-type and *lmd-3* mutants (n = 3 biological replicates, animals per condition per replicate ≥ 6, unpaired t-tests: ***$p < 0.001$). Scale bar, 10 μm. **j** Biochemical interaction of GFP-VHA-13 and mCherry-TLDc fragment demonstrated by co-IP and western blot of transgenic animal cell lysates. Co-transfected lysates were immunoprecipitated with GFP-Trap and immunoblotted for GFP and mCherry. Source data are provided as a Source Data file.

**Supplementary Fig. 12**

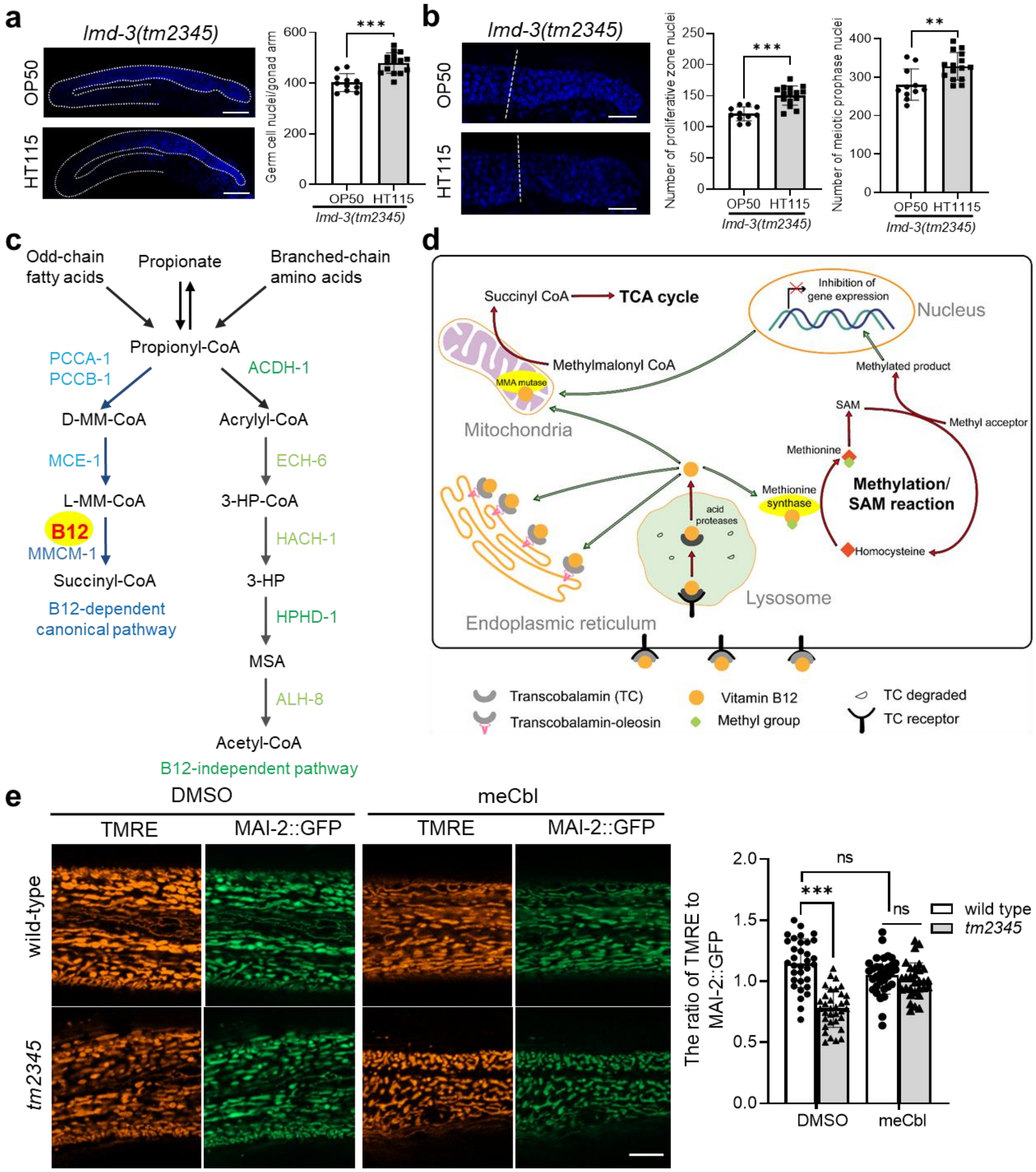

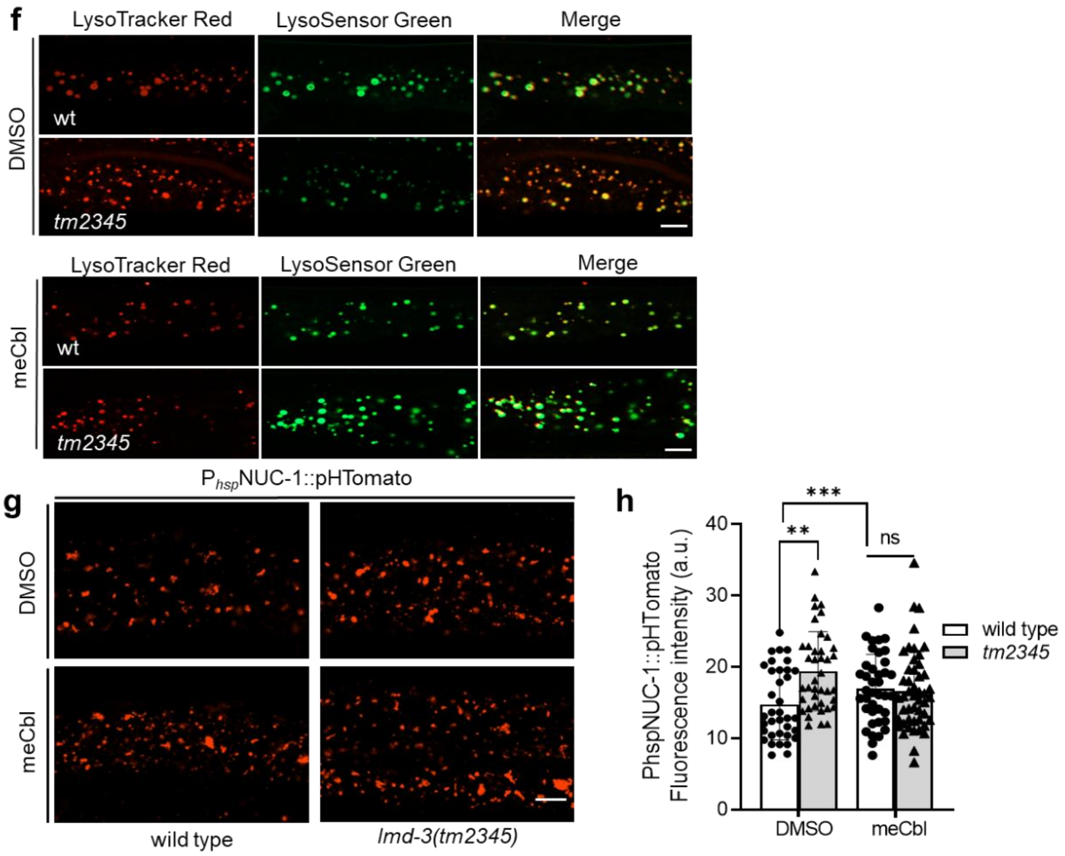


**Supplementary Fig. 12 Vitamin B12 restores reproductive function by alleviating cellular stress and improving organelle function. a** Exemplar fluorescence images and quantification of DAPI staining for total germline cells feeding with OP50 or HT115 in wild-type and *tm2345* mutants (n ≥ 11 for each group, unpaired t-tests: ***$p$ < 0.001). Scale bars, 50 μm. **b** exemplar fluorescence images and quantification of DAPI staining for germline stem cells and meiotic cells in wild-type and *tm2345* mutants (n ≥ 11 for each group, unpaired t-tests: **$p$ < 0.01, ***$p$ < 0.001). Scale bars, 20 μm. **c** Cartoon of branched-chain amino acids, propionic acid and odd-chain fatty acids metabolic pathways of in *C. elegans*. B12, vitamin B12; 3-HP, 3-hydroxypropionate; MM-CoA, methylmalonyl-CoA; MSA, malonic semialdehyde. **d** Model depicting the vitamin B12 uptake and metabolic roles. Cells internalize B12-bound transcobalamin (TC) via transcobalamin receptor (TCR)-mediated endocytosis and lysosomal processing.

Released B12 functions as adenosylcobalamin in the TCA cycle (propionyl-CoA to succinyl-CoA) and as methylcobalamin in cytosolic folate/methionine cycles. B12 deficiency disrupts DNA methylation and gene expression in *C. elegans*. SAM, S-Adenosyl methionine; TCA, tricarboxylic acid; Green arrows: transport; red arrows: metabolic pathways. **e** Confocal image and quantification of mitochondrial membrane potential in wild type and *tm2345* mutant with meCbl treatment ($n \geq 32$ for each group, unpaired t-tests: ***$p < 0.001$, ns, no significant differences). Scale bars, 10 μm. **f** Confocal fluorescence images of LSG/LTR co-stained intestines of meCbl-treated wild type and *lmd-3(tm2345)* animals at the D1 stage. Scale bars, 10 μm. **g, h** Exemplar fluorescence images (**g**) and quantification (**h**) of the heat-shock (hs) promoter driven NUC-1::pHTomato in the hypodermis at D1 adults ($n \geq 36$ for each group, unpaired t-tests: **$p < 0.01$, ***$p < 0.001$, ns, no significant differences). a.u., arbitrary units. Scale bar, 10 μm. Source data are provided as a Source Data file.

**Supplementary Table 1. Primers and oligos used in this study.** (See Excel spreadsheet uploaded separately)

**Supplementary Table 2. List of proteins identified from IP-MS** (See Excel spreadsheet uploaded separately)

**Supplementary Table 3. RNAi of LMD-3-interacting proteins for phenotypic analysis of reproductive capacity.** (See Excel spreadsheet uploaded separately)

**Supplementary Table 4. *C. elegans* strains used in this study.** (See Excel spreadsheet uploaded separately)
